# *foxQ2* marks fast-acting interneurons including dopaminergic neurons of *mushroom bodies* and *central complex* in the beetle *T. castaneum*

**DOI:** 10.64898/2026.05.11.724235

**Authors:** Ying Pang, Bastian Klußmann-Fricke, Doga Cedden, Jiajun Zhang, Johannes B. Schinko, Michalis Averof, Thomas Riemensperger, Gregor Bucher

## Abstract

The brain is one of the most complex animal organs and the development of the many different neuron types remains enigmatic. A set of brain-specific transcription factors is involved in brain patterning but their specific contributions mostly remain to be elucidated, including *foxQ2II*. This transcription factor is conserved in anterior neuroectodermal patterning of most animals while it has been lost from vertebrates. The contribution of *foxQ2II*-positive neurons to the adult brain has remained enigmatic. Here, we use our enhancer trap, immunostainings and our newly established beetle *brainbow* system to categorize *Tc-foxQ2II*-positive neurons into nine clusters with different projection patterns. All clusters contain neurons with the fast-activating neurotransmitters acetylcholine and glutamate while no *Tc-foxQ2II* positive neuron is GABA-ergic or serotonin-positive. Interestingly, we found that many dopaminergic neurons were *Tc-foxQ2II* positive and we homologize them with dopaminergic neurons of the PPL2c, PPM1 and PPL1 cluster described in the *Drosophila* brain. Our results show that *Tc-foxQ2II* marks subsets of fast-acting interneurons contributing to the higher order brain centers *mushroom bodies* and *central complex*. Taken together, our work expands the known functional range of *foxQ2* genes from sensory and neurosecretory cell specification to interneurons involved in the function of higher order brain centers.

## Introduction

The brain is built by a plethora of different neural cell types, which are interconnected in a very complex yet strictly ordered way. How this complexity develops and how the neural cell types are specified remains one of the major enigmas of developmental biology. Insects have served as an important model organism to understand basic principles of brain development because they have comparably simple brains and are experimentally more accessible than vertebrates. Especially the fruit fly *Drosophila melanogaster* has served as excellent model organism to pioneer molecular and genetic insights into nervous system development (Bellen et al., 2010). Contrasting limited knowledge about brain development, extensive research has revealed the principles of neural pattern formation and genetic signals of cell specification in the ventral nerve cord (Skeath and Thor, 2003; Urbach and Technau, 2004). In brief, the segmental neuroectoderm is subdivided by an orthogonal grid formed by the expression of columnar and segment polarity genes while Hox gene expression conveys regional identity. When the neural stem cells (neuroblasts, NBs) delaminate from the neuroectoderm, they inherit this spatial pattern of transcription factors from the neuroectoderm (e.g. different combinations of columnar, segment polarity and other genes). This “transcription factor cocktail” confers different identities to the delaminating NBs, which subsequently develop largely cell-autonomously into different neural lineages. On top of this spatial patterning, further neural cell diversification occurs e.g. by temporal patterning and lateral inhibition (Miyares and Lee, 2019; Skeath and Thor, 2003; Truman et al., 2010). In holometabolous insects, the embryonic phase of brain development leads to a simplified but functional larval brain. Postembryonic development adds to the scaffold provided by the larval brain and eventually leads to the formation of the adult brain (Hartenstein et al., 2008; Truman and Riddiford, 2023).

The insect brain is composed of three neuromeres, which stem from different neuroectodermal regions. The tritocerebrum and deutocerebrum derive from the intercalary and antennal segments, respectively. Due to their segmental origin, the basic principles of patterning these neuromeres are likely to be similar to the ventral nerve cord and indeed the expression of patterning genes in their NBs is similar (Urbach and Technau, 2003b; Urbach and Technau, 2004). The protocerebrum differs from the segmented ventral nerve cord in several respects. It contains crucial neuropils not found more posteriorly such as the *mushroom bodies,* the *central complex* and the *optic lobes*. Further, it is built by a much larger pool of NBs and in addition to type I neuroblasts, which build the VNC, the brain additionally contains some type II NBs. These type II NBs produce more offspring by producing intermediate neural progenitors, which themselves divide in a stem cell like fashion and contribute to the central complex (Bello et al., 2008; Boone and Doe, 2008; Boyan et al., 2010; Walsh and Doe, 2017). We and others have argued that the protocerebrum derives from the anterior non-segmental part of the neuroectoderm and evolutionary predated the segmentally organized ganglia. This lends an evolutionary explanation for pronounced differences with the VNC (Posnien et al., 2023; Scholtz and Edgecombe, 2006).

Molecular divergence reflects these differences but in contrast to the VNC, the genes and interactions involved in brain development are much less comprehensively studied (Kunz et al., 2012; Urbach and Technau, 2003b; Urbach et al., 2003). It is clear, however, that a different set of genes must be involved because the Hox genes and most columnar and segment polarity genes are not expressed in the anterior-most region of the embryo. Instead, a different set of highly conserved transcription factors is involved in anterior head and brain patterning in all animals (Gehring and Ikeo, 1999; Hirth et al., 2003; Leclère et al., 2016; Posnien et al., 2011; Posnien et al., 2023; Simeone et al., 1992; Sinigaglia et al., 2022; Tomer et al., 2010). The regulatory input by this set of uniquely anterior genes likely contributes to the development and specification of insect brain-specific structures such as the *central complex*, mushroom body and optic lobe neuropils (Posnien et al., 2023). However, the function of most of these genes in insect brain development remains unknown.

### Some *foxQ2* gene family transcription factors are involved in early anterior and neural patterning

Three foxQ2 genes are thought to have existed in the ancestors of all bilaterians. A complex pattern of losses led to a situation where the *foxQ2II* family is the only one retained in many protostomes (e.g. arthropods, nematodes) while it had been lost in tunicates and vertebrates including frogs, mice and fish (Gattoni et al., 2025b). Here, we focus on *foxQ2II,* which has been studied in many animals for its early anterior function and is the only paralog present in arthropods. In previous works in arthopods, this gene had been referred to as *foxQ2 (Tc-foxQ2* in *Tribolium)* and it corresponds to *Nematostella foxQ2a. foxQ2II* genes show an almost exclusive anterior expression in embryos of protostomes such as the fruit fly (Lee and Frasch, 2004), the red flour beetle (He et al., 2019; Kitzmann et al., 2017), spiders (Schacht et al., 2020) and annelids (He et al., 2019; Kitzmann et al., 2017; Marlow et al., 2014). Similar polar expression is found in deuterostomes such as the sea urchin and basal chordates while that paralogue has been lost in tunicates and vertebrates (Gattoni et al., 2025b; Yaguchi et al., 2008). The gene is expressed at the aboral pole in a sister clade to all bilaterian animals, the Cnidarians, indicating an ancestral function (Sinigaglia et al., 2013). Other *foxQ2* family members were found to be expressed in anterior neural calls in basal chordates and were shown to be required for retinal patterning in fish (Gattoni et al., 2025a; Gattoni et al., 2025b; Ogawa et al., 2021). Involvement in sensory cell development has been suggested for the *foxQ2d* paralog of a Cnidarian (Busengdal and Rentzsch, 2017).

While some members of the *foxQ2* family have been related to neurosecretory and sensory cell development, information on the neural types marked by *foxQ2II* has remained scarce in animals and absent from arthropods. This includes the fly ortholog *fd102C* for which functional data has been lacking (Lee and Frasch, 2004). *Drosophila* has a rather derived mode of head development: During embryogenesis, the head and brain neuroectoderm are involuted outside-into the thorax leading to a highly diverged and reduced structure, which has been difficult to analyze. *Drosophila* is also derived within insects with respect to its comparably small number of brain neuroblasts (Urbach and Technau, 2003a), and its simplified larval brain, which for instance lacks a *central complex* neuropil (Koniszewski et al., 2016; Pfeiffer and Homberg, 2014). In order to study the genetic control of brain development in a more insect-typical situation, the red flour beetle *Tribolium castaneum* has been established as complementary model system (Posnien et al., 2010). The extensive toolkit available in this species includes transgenesis, genome editing, a strong RNAi response and genome-wide RNAi resources, which together allow for in depth functional studies (Klingler and Bucher, 2022). For instance, new insights have been gained into the heterochrony of the *central complex* development and a diverging number of type II neuroblasts was found compared to flies (Farnworth et al., 2020; Rethemeier et al., 2025).

In *Tribolium*, the role of *foxQ2II* has been well characterized with respect to embryonic brain development. In the embryo, *Tc-foxQ2II* is co-expressed with *six3/optix* and plays a key role at the top of the anterior gene regulatory network (aGRN) (Kitzmann et al., 2017), where it interacts with other brain-specific transcription factors, such as *Tc-six3/optix*, *Tc-six4*, *Tc-chx/vsx*, *Tc-nkx2.1/scro*, *Tc-ey*, *Tc-rx* and *Tc-fez1* (Kitzmann et al., 2017). Further, *Tc-foxQ2II* is expressed in several embryonic neuroblasts, which have at least four different molecular identities based on co-expression with other genes. Probably, both type I and type II NBs are marked (He et al., 2019). RNAi knock-down of *Tc-foxQ2II* prohibits the split of the primary brain commissure into several fascicles, from which parts of the *central complex* develop. This leads to severe disturbance of the *central complex* and secondarily to the aberrant arrangement of the mushroom body (He et al., 2019; Kitzmann et al., 2017). To identify the neural cells marked by *Tc-foxQ2II,* an enhancer trap line was generated by genome editing. This revealed that in the larval brain, *Tc-foxQ2II* enhancer trap marks at least three clusters of neurons but no glia. Some neurons project into the larval *central body* among other structures (He et al., 2019).

In contrast to embryonic development, the role of *foxQ2II* during postembryonic arthropod development remains unknown. Here, we characterize the *Tc-foxQ2II* marked neurons in the adult brain. We identify and characterize nine neuron clusters and we established the brainbow system in *Tribolium* to confirm projections of these cells by sparse labelling. We find that many *Tc-foxQ2II* marked neurons project into the *mushroom bodies* or the *central complex*. Further, all *Tc-foxQ2II* positive neurons are positive for the fast-acting neurotransmitters acetylcholine and glutamate while we find no coexpression with GABA or serotonin. A subset is dopamine and octopamine positive including a large part (∼71.2%) of the adult dopaminergic neurons, which probably correspond to fly neurons from the PPL2c, PPM1 and PPL1 clusters. Taken together, these results suggest an important role of *foxQ2II* in the specification of interneurons whereas in other animal clades, sensory and neuroendocrine cell types have been linked to *foxQ2*. Further, we find that *Tc-foxQ2II* positive neurons contribute to the higher order brain centers including both, columnar and tangential neurons of the *central complex*.

## Results

### *Tc-foxQ2II* positive cells of the adult brain are organized into nine clusters

*Tc-foxQ2II* positive neurons contribute to the *central complex* both in the larval and the adult brain (He et al., 2019). While the larval projection pattern has been well described in *Tribolium*, detailed information on the adult brain has been lacking in any insect. To close this knowledge gap, we used our established transgenic *foxQ2II* imaging line (*foxQ2-5’-line*), which marks *Tc-foxQ2II* positive cells with EGFP and Cre (He et al., 2019). To check for the degree of overlap of EGFP and Tc-foxQ2II expression in adult brains, we performed double immunohistochemistry with anti-FoxQ2 and anti-GFP antibodies (He et al., 2019). We detected that 85.1% of cells expressing Tc-foxQ2II also expressed EGFP, while 79.0% of EGFP-expressing cells overlapped with Tc-foxQ2II (n= 3; Supplementary Fig. 1 A-A’’ and B). We concluded that the expression patterns overlap sufficiently to use EGFP expression of the *foxQ2-5’-line* as proxy for studying match *Tc-foxQ2II* positive cells.

We found nine *Tc-foxQ2II* positive cell clusters in the adult brain based on cell body location and their main projections (Fig. 1 A-D; Fig. 2 A,B). One additional cluster (anterior cluster, AC; white star / yellow color in Fig. 1 A’, B’, and Fig. 2) was not marked by the Tc-foxQ2II antibody (supplementary Fig. 2) and was therefore not considered further in this study. Three primary clusters had been assigned by He for the embryonic and larval brain (He et al., 2019): The median cluster (MC), the lateral cluster (LC) and the posterior cluster (PC). We named the nine clusters in the adult brain in accordance with this nomenclature. One brain was used for analysis and 3D reconstructions (Fig. 1 and subsequent figures) while the statements were confirmed in five additional brains (the original image stacks are available for download and further processing at Gro.data (https://doi.org/10.25625/FWBHWP) and AVI formats can be viewed on Figshare (doi: 10.6084/m9.figshare.30531422).

**Figure 1.**
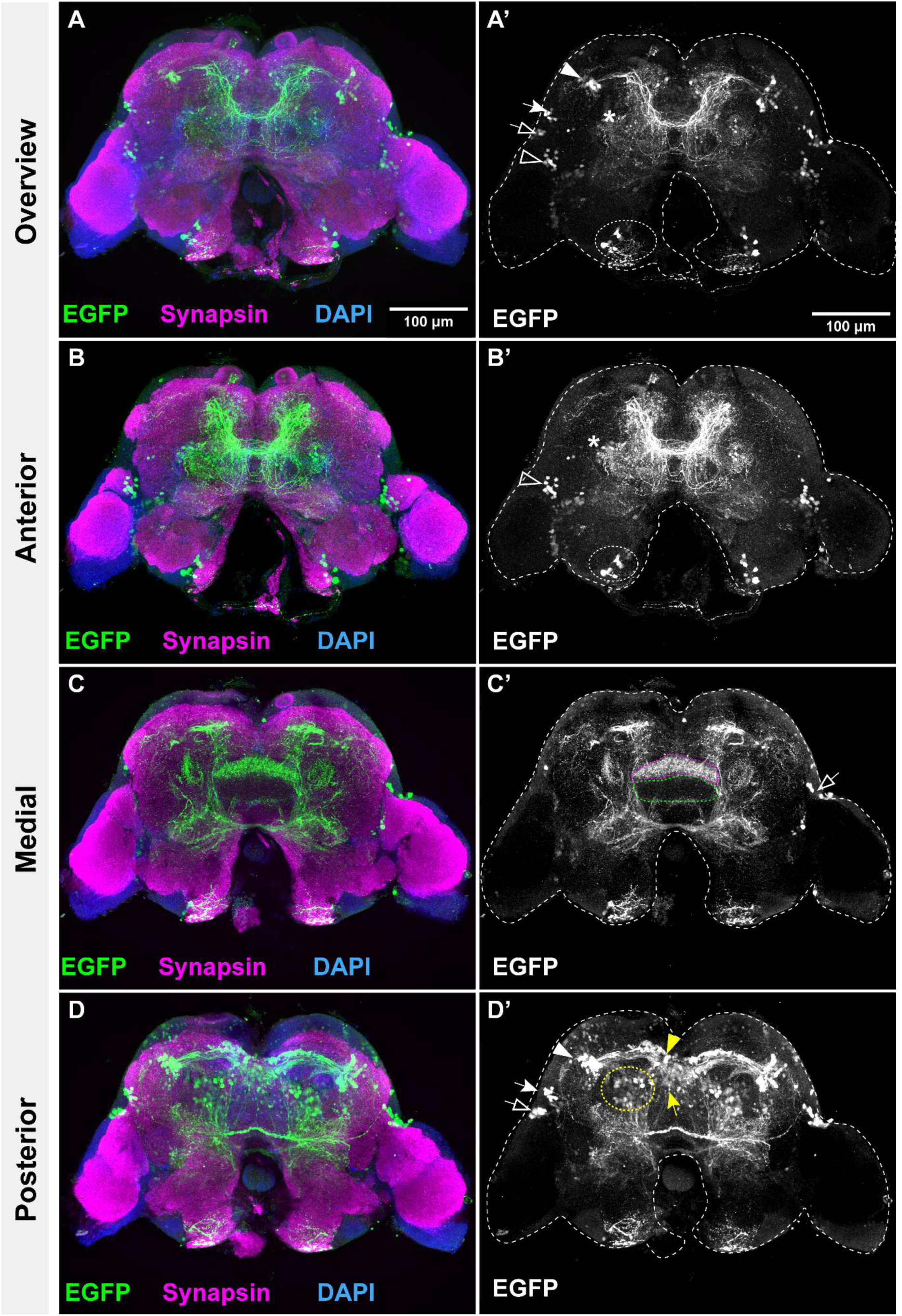
Overview on the *Tc-foxQ2II* positive cells. Neuronal projection patterns were identified using immunohistochemistry against EGFP in the adult brain of *foxQ2-5’-line.* (A-D) Different parts of the adult brain stained for EGFP (green), Synapsin (magenta), and DAPI (blue). (A’-D’) The EGFP channel is shown with the brain outline indicated by dashed lines. (A-A’) Overview of a projection of the entire brain; (B-B’) anterior subsection; (C-C’) medial subsection; (D-D’) posterior subsection. White arrowhead: Lateral Cluster 1; white arrow: Lateral Cluster 2; open arrowhead: Lateral Cluster 3; open arrow: Lateral Cluster 4; yellow arrowhead: Median Cluster 1; yellow arrow: Median Cluster 2; yellow dashed circle: Posterior Cluster; white dashed circle: Tritocerebrum Cluster; white star: Anterior Cluster. Scale bar: 100 μm.

**Figure 2.**
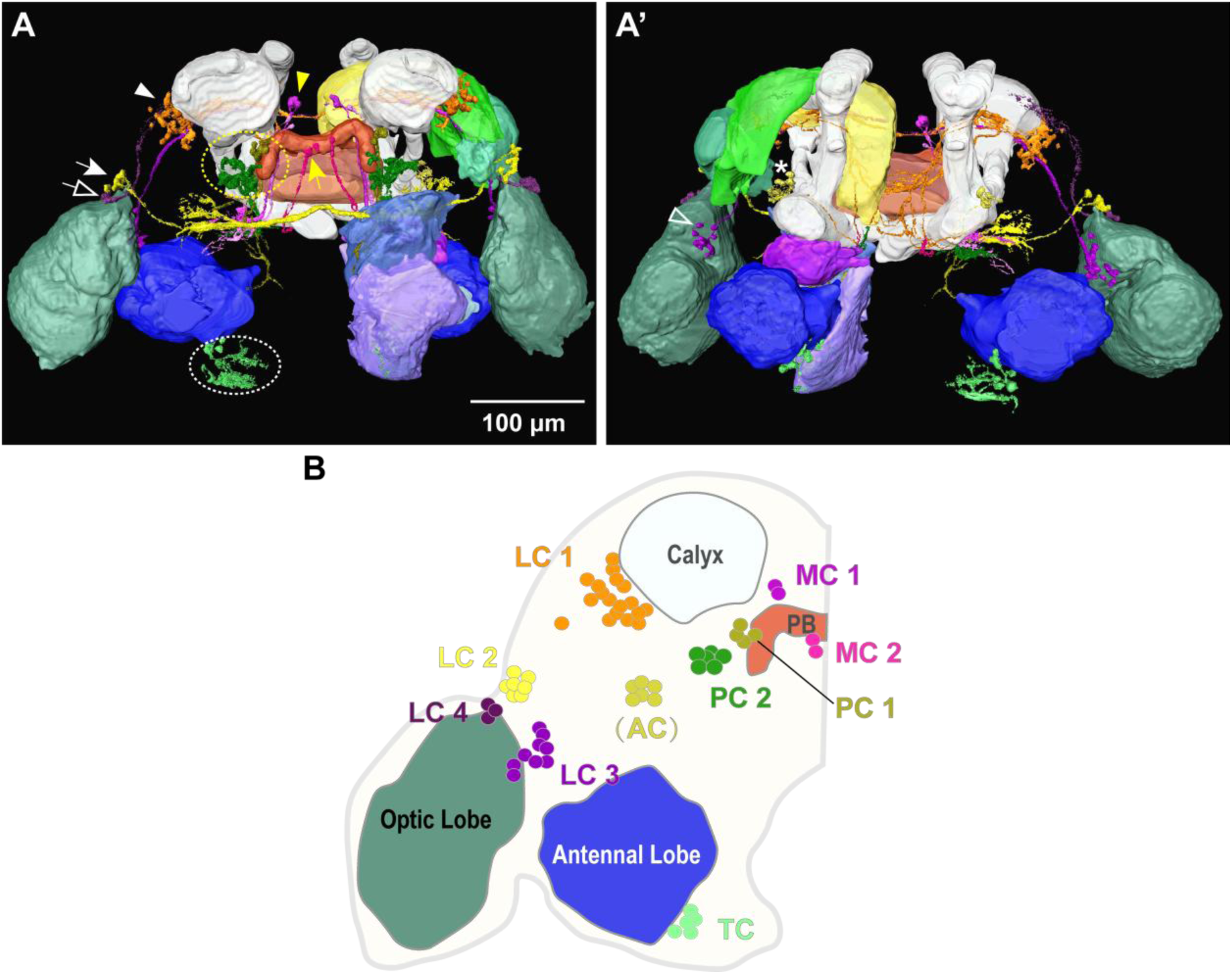
Overview on the reconstruction of the nine *Tc-foxQ2II* positive cell clusters. (A) 3D reconstruction of major neuropils and EGFP-positive clusters of neurons based the brain shown in Fig. 1. Posterior view in (A) and anterior view in (A’). The cluster AC (white star in A’) was not analyzed further as it was not Tc-foxQ2II positive. White circle: Tritocerebral cluster; yellow circle: Posterior cluster 1 and 2. (B) Schematic overview of the cell body locations for the ten clusters shown in the same colors as in A/A’ and subsequent figures. White arrowhead: Lateral Cluster 1 (orange); white arrow: Lateral Cluster 2 (yellow); open arrowhead: Lateral Cluster 3 (purple); open arrow: Lateral Cluster 4 (dark purple); yellow arrowhead: Median Cluster 1 (magenta); yellow arrow: Median Cluster 2 (pink); yellow dashed circle: Posterior Cluster (PC1 is olive green and PC2 is vibrant green); white dashed circle: Tritocerebrum Cluster (light green and white broken circle in A); white star: Anterior Cluster (grass yellow). Scale bar: 100 μm.

#### Ipsilateral projection neurons (MC1 and MC2) located near the central complex

The neuropils reconstructed based on synapsin staining and their abbreviations are shown in Fig. 3A. The cell bodies of MC1 (magenta in Fig. 3B,B’) are located above the protocerebral bridge (PB) and project two long tracts through the PB (we found no projections into the PB) towards the posterior slope (VMCpo/PS). According to the standard neuropil compartmentalization (Farnworth et al., 2022), MC1 projections traverse the inferior protocerebrum, posterior domain (IPp) and the posterior slope (VMCpo/PS), with partial projections ultimately reaching the boundary between the VMCpo/PS and the lateral accessory lobe (LAL) and arborizing within the LAL. Compared to MC1, the cell bodies of MC2 (Fig. 3 C-C’) are positioned closer to the midline, residing in the central depression of the PB. Their projections do not cross the midline but extend a tract towards the VMCpo/PS and LAL.

**Figure 3.**
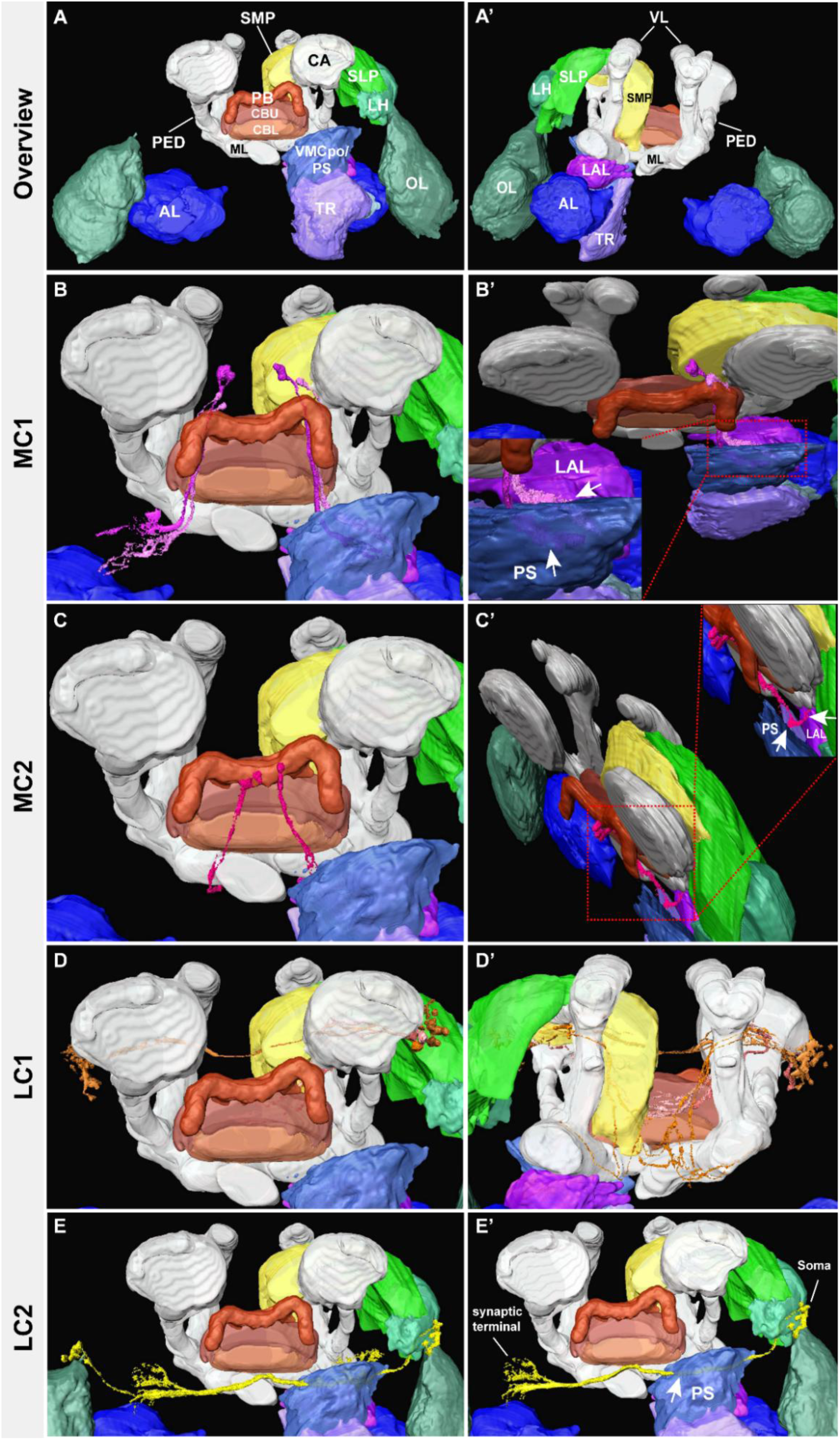
The morphology and distribution of EGFP-positive clusters. (A-A’) Posterior view and anterior views of the brain, showing major neuropils. (B-B’) Median cluster 1 (MC1). (B) Posterior view. (B’) Lateral view. The region enclosed by the red dashed square is magnified in the bottom-left corner, with a white arrow indicating the specific projection area. (C-C’) Median cluster 2 (MC2). (C) Posterior view. (C’) Lateral view. The central complex (CX) region, enclosed by a red dashed square, is magnified in the top-right corner. A white arrow indicates projections targeting the VMCpo/PS and LAL. (D-D’) Lateral cluster 1 (LC1). (D) Posterior view. (D’) Anterior view. (E-E’) Posterior view of lateral cluster 2 (LC2). In (E’), a white arrow marks the VMCpo/PS region traversed by LC2. Abbreviations: AL: Antennal lobe; CB: Central body (CBU: Upper unit of the central body; CBL: Lower unit of the central body); LH: Lateral horn; MB: Mushroom body (CA: Calyx; ML: Medial lobe; PED: Peduncle; VL: Vertical lobe); OL: Optic lobe; PB: Protocerebral bridge; SLP: Superior lateral protocerebrum; SMP: Superior medial protocerebrum; TR: Tritocerebrum; VMCpo/PS: Ventromedial cerebrum postcommissural domain / Posterior slope.

The location of the cell bodies of MC1 neurons and their projection ventrally along the midline between the PB and the CB is similar to dopaminergic PPM1 cluster neuron from *Drosophila melanogaster* (Supplementary Fig. 3 A-A’) (Hartenstein et al., 2017; Mao and Davis, 2009), typed as PPM1205 in the *Drosophila* electron microscopy dataset (Schlegel et al., 2024). MC2 neurons may equally correspond to the dopaminergic neurons of the same PPM1 cluster in adult *Drosophila* and may be assigned to the neuron PPM1204 in the *Drosophila* electron microscopy dataset (Schlegel et al., 2024).

#### The lateral cluster 1 comprises tangential neurons innervating the mushroom body and the central complex

The projections and location of cell bodies allowed classifying lateral clusters (LC) into four subclusters: LC1-4. The cell bodies of LC1 (Fig. 3 D-D’) were located adjacent to the calyx (CA; MBs are white in Fig. 3), with their projections passing through the CA and the superior medial protocerebrum (SMP; yellow). Two different projections seem to emanate from LC1. The projection of LC1a (orange in Fig. 5 A-A’) has two primary branches: one first circumvents the vertical lobe (VL, white) of the mushroom body (MB, white), then projects through the crepine toward the ventral region, and eventually arborizes in the ML (Fig. 5 A-A’). The other, LC1b (light coral in Fig. 5 B-B’), directly forms arborizations within the CB (white arrowhead in Fig. 5 A’). It does not circumvent the VL but projects through the superior lateral and medial protocerebrum directly between the VL and the peduncle (PED, white) of the MB, arborizing in the anterior surface of the CB (Fig. 5 B-B’).

Regarding the cell body location and the projections mainly to the mushroom body and the central complex (Supplementary Fig. 3 C-C’), LC1 cluster neurons appear similar to dopaminergic PPL1 cluster neurons in the *Drosophila* brain (see below for further data) (Mao and Davis, 2011; Hartenstein et al., 2017). The LC1a subcluster appears to innervate mainly the medial lobes of the mushroom bodies, similar to PPL101 dopaminergic neurons in *Drosophila* (Fig. 5 A’). Using immunohistochemistry, we confirmed dopamine expression in LC1a neurons that was expected for PPL1 neurons in *Drosophila* (see below). From the electron microscopy datasets (Scheffer et al., 2020; Schlegel et al., 2024) it appears that one at least one dopaminergic PPL1 neuron, PPL108 (Supplementary Fig. 3) displays an interconnection between the horizontal mushroom body lobes and the noduli of the central complex.

#### Lateral cluster 2-4 cells lie adjacent to the optic lobes and exhibit divergent projections

In LC2 (Fig. 3 E-E’), the cell bodies are located near the lateral horn (LH; mint green), with near-horizontal projections traversing the VMCpo/PS and ultimately projecting toward the VMCpo/PS of the opposite hemisphere. The cell bodies of LC3 (Fig. 4 A-A’) are located close to the optic lobe (OL; dark green), traverse the LH and the SLP, and arborize within the SMP (Fig. 4 A-A’). Based on the projection into the main neuropils and comparison with neurons in the *Drosophila* brain, LC3 is tentatively identified as a neuron similar to PPL2ab (Supplementary Fig. 3 E-E’) (Farnworth et al., 2022; Mao and Davis, 2009). LC4 (Fig. 4 B-B’) projects from the OL, traverses the LH and arborizes in the SLP (indicated by white arrows in Fig. 4 B’). Taken together, the LC2-3 neurons have their cell bodies close to the optic lobes and project to the contralateral hemisphere without overt connectivity to MB or CX.

**Figure 4.**
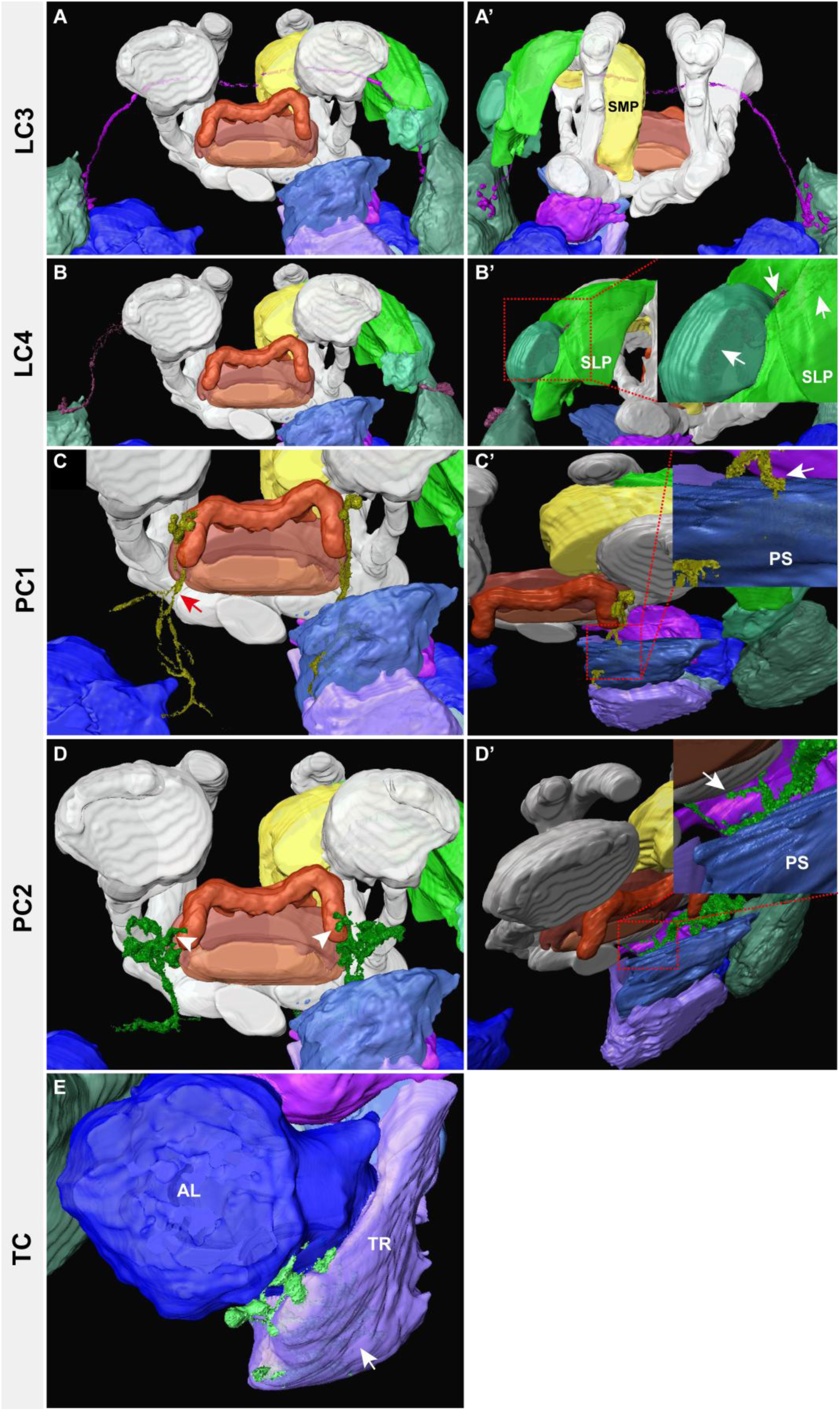
The morphology and distribution of EGFP-positive clusters in the brain of the *foxQ2-5’ line*. (A-A’) Lateral cluster 3 (LC3). (A) Posterior view. (A’) Anterior view. (B-B’) Lateral cluster 4 (LC4). (B) Posterior view. (B’) Lateral view. A white arrow indicates LC4 arborization in the LH and SLP. (C-C’) Posterior cluster 1 (PC1). (C) Posterior view. (C’) Lateral view. The region enclosed by the red dashed square is magnified in the top-right corner. A white arrow shows where PC1 enters the VMCpo/PS. (D-D’) Posterior cluster 2 (PC2). (D) Posterior view. (D’) Lateral view. The projection region, enclosed by a red dashed square, is magnified in the top-right corner. A white arrow points to the arborization site. (E) Anterior view of tritocerebrum cluster (TC). A white arrow indicates its arborization zone. Abbreviations: AL: Antennal lobe; CB: Central body (CBU: Upper unit of the central body; CBL: Lower unit of the central body); LH: Lateral horn; MB: Mushroom body (CA: Calyx; ML: Medial lobe; PED: Peduncle; VL: Vertical lobe); OL: Optic lobe; PB: Protocerebral bridge; SLP: Superior lateral protocerebrum; SMP: Superior medial protocerebrum; TR: Tritocerebrum; VMCpo/PS: Ventromedial cerebrum postcommissural domain / Posterior slope.

Note that due to the common origin of the four major projections from one large lateral cell cluster and due to the lack of single cell resolution in this analysis, the described assignments into LC1-4 subclusters remained tentative. We established and applied sparse marking of cells using the Brainbow system and added neurotransmitter analyses to confirm the interpretation (see below).

#### Posterior cluster cell bodies are located close to the PB but do not project into the central complex

The PC is subdivided into two subclusters based on the locations of cell bodies and their projections. The cell bodies of PC1 (Fig. 4 C-C’) are located adjacent to the lateral PB and project to the VMCpo/PS, forming Y-shaped arborizations in the VMCpo/PS of the central brain (red arrow in Fig. 4 C-C’). PC2 cell bodies are distributed in the inferior protocerebrum, central domain (IPc), located beneath the CA, and project to the VMCpo/PS and the LAL (Fig. 4 D-D’; white arrowhead marks cell bodies in D). Based on these characteristics, we hypothesize that the PC1 neurons may be homologous to the descending neurons (DNs) of *Drosophila* (Supplementary Fig. 3 F-F’) (Namiki et al., 2018).

#### The Tritocerebrum Cluster

In the adult brain we found a cluster not described in He et al. and we named it TC (tritocerebrum cluster). The cell bodies of the TC are located between the antennal lobe (AL; blue) and the tritocerebrum (TR; light purple; Fig. 4E and white dashed circles in Fig. 1 A’-B’ and Fig. 2 A) and some projections are found within the TC (white arrow in Fig. 4 E). To check, whether the TC emerges only during postembryonic development or had been overlooked previously, we did stainings in the L1 larval brain. Indeed, we found cells, which prefigure TC at that stage (white arrows in Supplementary Fig. 4).

In addition to the above described clusters, there were also cells whose projections could not be visualized or clearly tracked due to low signal (white arrows in Supplementary Fig. 5). In the absence of identifiable projections, we refrained from homologizing these cells.

### Establishing the brainbow system in *Tribolium castaneum* for sparse marking of neurons

Due to the large number of marked neurons in the *foxQ2-5’ line* and the resulting overlapping projection patterns, it was difficult to unequivocally reconstruct the projection patterns of certain clusters (e.g. LC1). In order to test our hypotheses by an independent approach, we used sparse labelling of *Tc-foxQ2II* positive neurons by establishing and using the Brainbow system in *Tribolium* (Livet et al., 2007). This was facilitated by the fact that the *foxQ2-5’-line* has a bicistronic transcription unit that codes for both, EGFP and the Cre-recombinase (Supplementary Fig. 6) (He et al., 2019). We generated a transgenic line with the Brainbow 1.1 M construct, which was modified in two ways: (a) the Brainbow cassette was placed under the *Tribolium* ubiquitous promoter EFA (Sarrazin et al., 2012), and (b) the first fluorescent protein (Kusabira) was mutated such that no fluorescence is detectable in the absence of Cre-mediated recombination (see Methods). By crossing this *EFA-Brainbow line* with the foxQ2-5’ line, membrane-bound mCherry, mEYFP or mCerulean are expected to be expressed stochastically in the *foxQ2II* neurons (see Supplementary Figure 6A). We could then observe the neurons expressing mCherry using an antibody raised against RFP (EYFP, Cerulean and the EGFP expressed by the foxQ2-5’ transgene could not be distinguished by immunostaining, as these proteins have closely related sequences). To test the system, we confirmed that mCherry expression from the *EFA-Brainbow line* was dependent on the presence of the Cre construct at 23 and 32°C (Supplementary Fig. 6 C and Supplementary Fig. 7). As the Cre recombinase activity increases with temperature (Buchholz et al., 1996), we tested the number of mCherry marked cells at two different temperatures, which reflect the upper and lower limits of the optimal living temperature of *Tribolium* (32 °C and 23 °C). In 20 brains grown at 23 °C, a total of 79 cells were marked (Supplementary Fig. 7). Some clusters were frequently marked with several cells, while others were marked much more rarely. The six brains grown at 32 °C showed no clear increase in the number of marked cells (Supplementary Fig. 7). As a comparably large number of cells were marked in each brain, the projections of the neurons of interest often converged with those of other neurons, which made unequivocal reconstruction of the entire projection pattern of the neurons impossible. In summary, the system allows for sparse random marking of cells but does not provide single cell resolution.

### Confirming cluster assignments using the *Tribolium brainbow* system

After mating the *foxQ2-5’-line* with the *EFA-Brainbow line*, we collected brains from the offspring (grown at 23°C) and stained for mCherry. From 20 brains we reconstructed in total 79 single marked neurons or small neural clusters. For conservative analysis, we selected cells that had no or comparably few marked neighbors and reconstructed only the main projections as far as they were unequivocal. We were able to confirm all clusters described above except for LC3 (Fig. 6).

Two subclusters had tentatively been assigned to LC1 (Fig. 5) and we used sparse Brainbow labeling to further test this hypothesis. We found that 10 cells from 9 brains reflected the LC1 projection pattern (Supplementary Fig. 8). After segmentation of these cells, we confirmed the presence of two subclusters LC1a and LC1b based on the fact that some neurons passed anterior of the vertical lobe of the MB (LC1a, Fig. 5A’) while others projected posterior to it into the central brain (LC1b, Fig. 5B’). Each was confirmed by 5 neurons, respectively (Supplementary Fig. 8).

**Figure 5.**
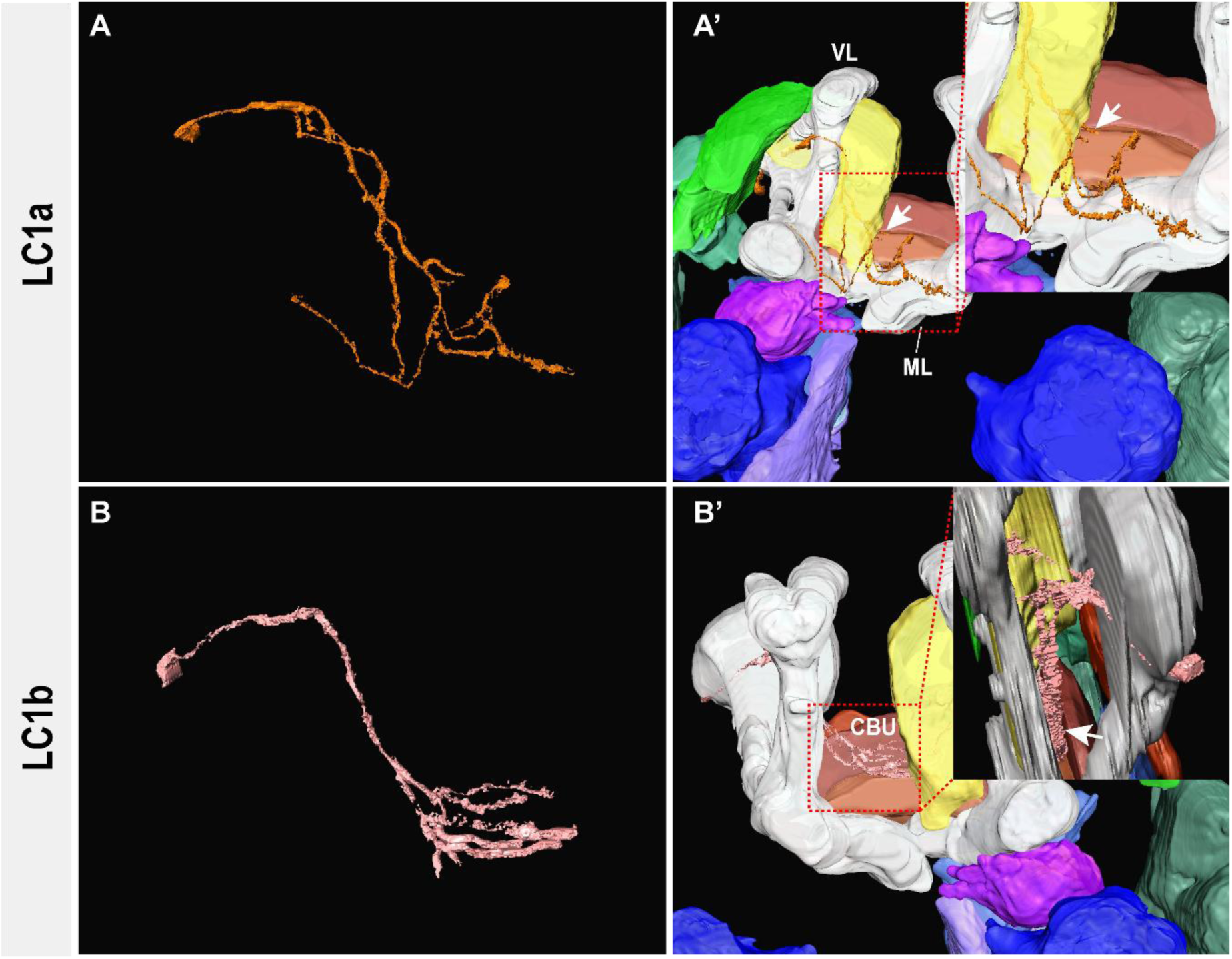
Classification of Lateral Cluster 1 based on distinct projection patterns. (A-A’) Morphology and projection pattern of LC1a, shown without (A) and with (A’) brain compartments. The region enclosed by the red dashed square shows where LC1a passes near the central body (CB) before crossing the midline. A magnified view of this region (top-right corner) shows the LC1a projection adjacent to the CB, indicated by a white arrow. (B-B’) LC1b projections. The boxed region in (B’) shows that LC1b projects toward the CB. A magnified lateral view (top-right) details the contact point (white arrow). Abbreviations: VL: the vertical lobe; ML: the medial lobe; CBU: the upper unit of the central body.

**Figure 6.**
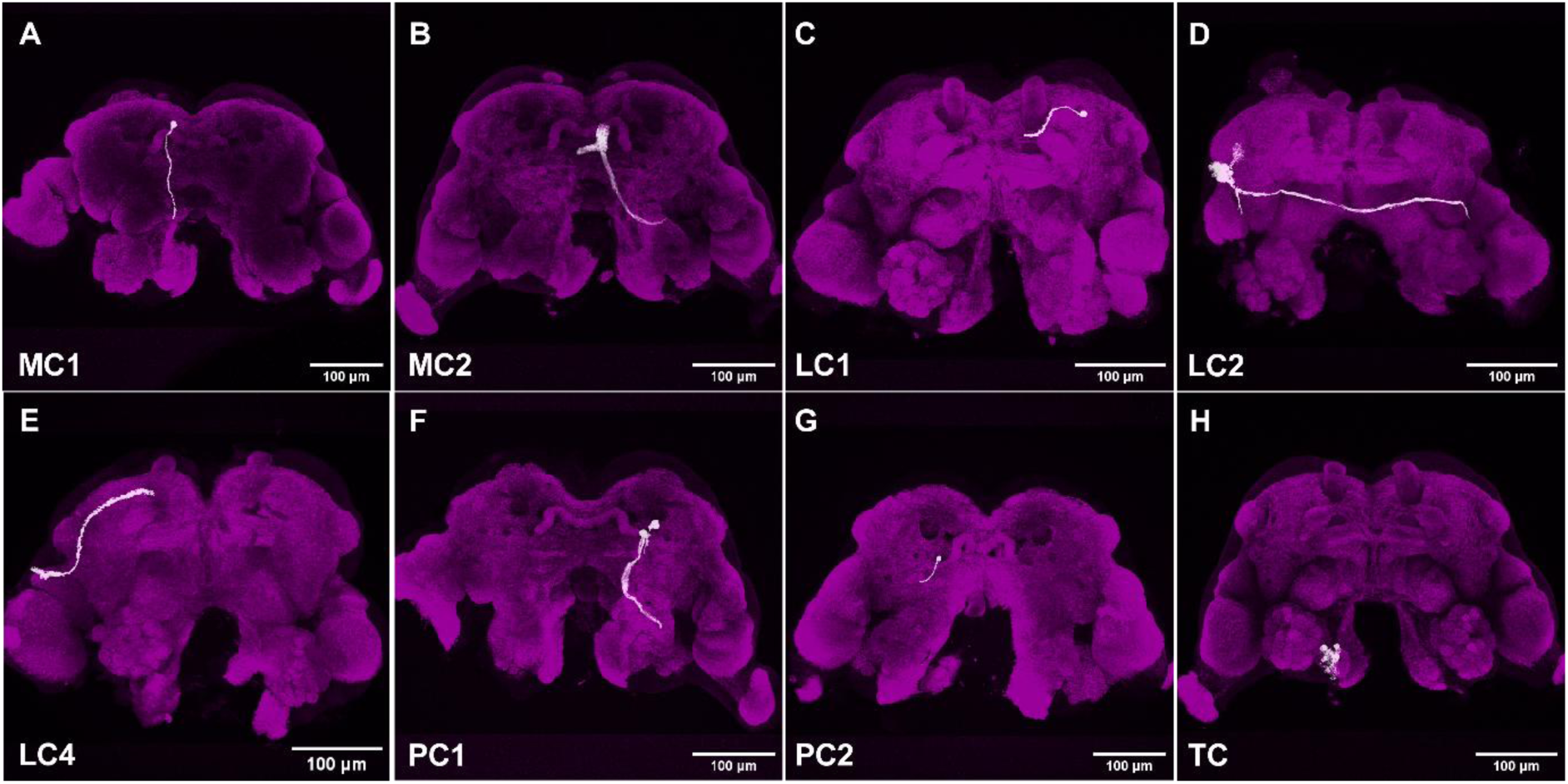
Eight clusters detected in the *foxQ2-5’– line* were also sparsely labelled by Brainbow. (A) MC1. (B) MC2. (C) LC1. (D) LC2. (E) LC4. (F) PC1. (G) PC2. (H) TC. All panels show the brain of the *foxQ2-5’- line*. Magenta represents synapsin. Scale bar: 100 μm.

### Embryonic expression of *Tc-foxQ2II* is not maintained in all daughter cells

The *foxQ2-5’-line* is an enhancer trap reflecting the current state of expression of *Tc-foxQ2*. In brainbow, in contrast, expression continues permanently once turned on by recombination. Therefore, discrepancy in the patterns can be observed if cells had turned on brainbow e.g. during embryonic *Tc-foxQ2II* expression but subsequently ceased to express Tc-foxQ2. Such cases would be detectable by observing mCherry projections in the adult brain, which were not found in the *foxQ2-5’-line*. Indeed, we observed Brainbow marked neurons in the adult brain, which had not been observed in the *foxQ2-5’-line* (Supplementary Fig. 9 and 10). Notably, columnar neurons were observed among these cells, connecting distinct neuropils of the central complex (CX) and other brain structures through four prominent tracts (termed the WXYZ tracts) (Supplementary Fig. 9 A). This marking was dependent on Cre from the *foxQ2-5’-line* because the *EFA-Brainbow line* alone showed no mCherry signal (Supplementary Fig. 6 C). We conclude that these cells had expressed *EGFP/Cre* for some time during development but had shut down expression at later stages. The marking of columnar neurons is in line with the embryonic expression of *Tc-foxQ2II* in Type II neural lineages (He et al., 2019). In summary, during development, *Tc-foxQ2II* expression seems to be shut down in subsets of neurons. It is noteworthy that some columnar neurons stem from early *Tc-foxQ2II* positive cells while ongoing *Tc-foxQ2II* expression marks no columnar cells but some tangential cells of the CX (among other types of cells).

### LC1a interneurons innervate the *mushroom body* and the *central complex*

In our imaging line, we found plenty arborizations within the MBs and the CX (Fig. 7; arrows in Fig. A’ and arrowhead in Fig. B’ and single optical sections cutting through these neuropils in Fig. 7A’’ and B’’). Specifically, LC1 seems to comprise cells strongly innervating the MB and/or CX. The projections of LC1a (Fig. 5A) appear to arborize in the mushroom body medial lobes similar to the projection pattern of the LC1a neurons from the dopaminergic PPL1 cluster in adult *Drosophila* that have been identified as PPL101 in the electron microscopy dataset (Scheffer et al., 2020; Schlegel et al., 2024) (Fig. 5 A’).

**Figure 7.**
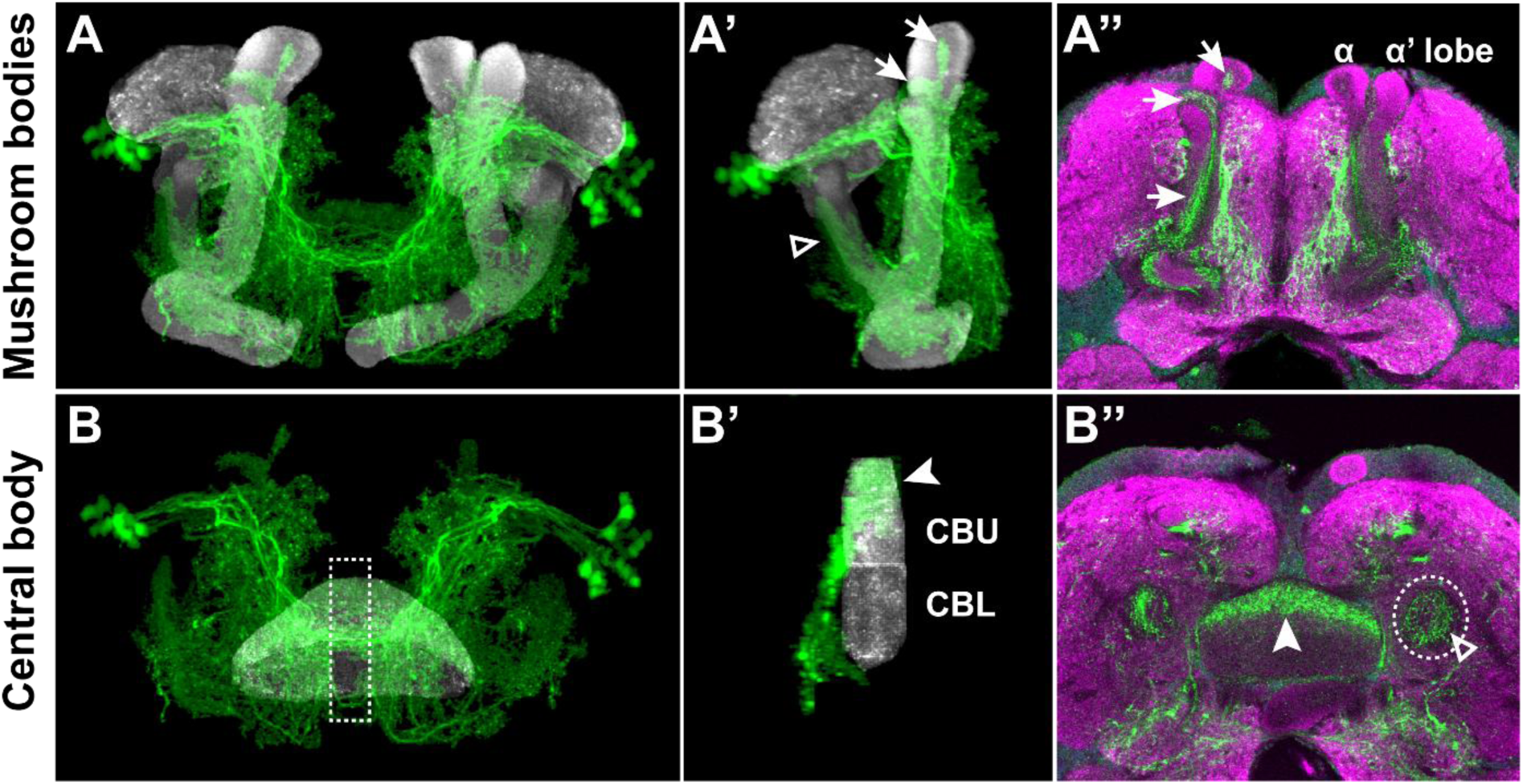
EGFP-positive cells arborize in the mushroom body and the central complex. (A-A’’) Arborizations of EGFP-positive cells in the mushroom body (MB). (A) Overview of 3D reconstruction of the MB and EGFP-positive projections, generated using VVDviewer. (A’) Hemi-mushroom body with EGFP-positive projections. (A”) Immunohistochemistry result showing the MB and EGFP-positive projections. White arrows in (A’) and (A’’) indicate EGFP-positive projections arborizing in the α and α’ lobe of the MB. (B-B’’) Arborizations of EGFP-positive cells in the central body (CB). (B) Overview of the 3D reconstruction of the CB with EGFP-positive projections. (B’) Side profile view of the area marked by the white dashed rectangle in (B). (B’’) Immunohistochemistry result showing the CB and EGFP-positive projections. White arrowheads in (B’) and (B’’) indicate EGFP-positive projections arborizing in the upper unit of the central body (CBU). Open arrowheads in (A’) and (B’’) indicate the EGFP-positive projections arborizing in the peduncle (PED). The White dashed circle in (B’’) outlines the PED. Green, EGFP; magenta, synapsin; and white, neuropils (MB or CB).

We wondered, whether single neurons would innervate both, MB and CX but the resolution of our standard LSM images did not allow separating the neurites clearly. To characterize the projections of these neurons more in detail, we performed STED high resolution microscopy of that region (Willig et al., 2006) and performed a more precise reconstruction of neurons belonging to the two subclusters of LC1 (Fig. 8A-B and Supplementary Fig. 18). LC1 neurons arborize in both the upper unit of the central body (CBU) and the medial lobe of the mushroom body (Fig. 8 A and Supplementary Fig. 18). In the CBU, they arborize extensively (yellow in Fig. 8C-F) while the tracts continue to the contralateral side meeting the other LC1 tracts at the midline (white in Fig. 8C-F). While the resolution of the STED images was clearly better (Supplementary Fig. 19) it did not suffice to unequivocally show or rule out that there are LC1 cluster neurons, which innervate both, MB and CX. This hypothesis will have to to be tested by enhanced tools for single cell marking (as opposed to sparse marking of our brainbow system). We note that there are EGFP-marked projections also in other parts of the MBs and the crepine (e.g. Fig. 7A’’ and B’’). Due to the dense labelling of *Tc-foxQ2II* positive neurons these could not be assigned unequivocally to certain neurons but they all seemed to be dopaminergic.

**Figure 8.**
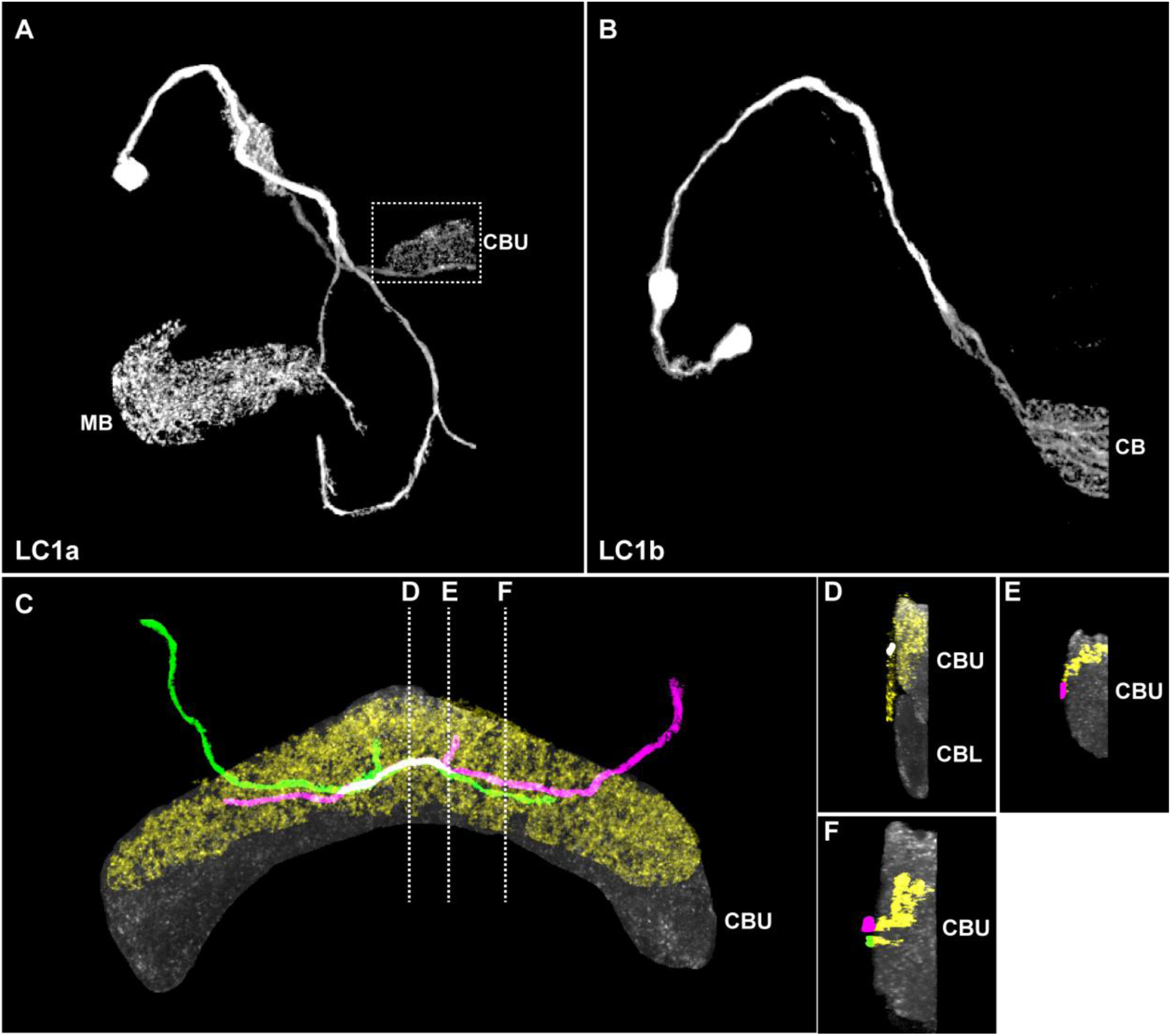
Reconstruction of STED-imaging of LC1a connection of mushroom body with the central complex in the *foxQ2-5’-line* brain. (A) LC1a seems to have two main branches: one innervates the mushroom body (MB), and the other innervates the upper unit of the central body (CBU). (B) LC1b arborizes only within the central body (CB). (C) A higher-magnification view of the projections in the CBU (region partially outlined by the white dashed rectangle in A). (D-F) Transverse sections through distinct domains of the central body. (D) Median section. (E) A lateral section showing a single branch projecting into the CB. (F) A section where two distinct tracts are visible. Green and magenta represent the main projections from different sides, which are in close proximity around the midline (white). Yellow indicates arborizations within the CBU. CBU is shown in grey. CBL: lower unit of the central body. The STED dataset used in this panel was acquired by Ying.

### Neurotransmitter content of *Tc-foxQ2II* positive cells

To better characterize the neuron subtypes within *Tc-foxQ2II* positive clusters, we examined the expression of six major neurotransmitters (Lacin et al. 2019). Neither GABA nor serotonin (5-HT) were expressed in neurons marked by EGFP (Table 1 and Supplementary Fig. 11 and 12). Most adult *Tc-foxQ2II* neurons were positive for the fast-activating neurotransmitters acetylcholine (visualized by immunohistochemistry targeting Choline Acetyltransferase (ChAT)) and Glutamate (Supplementary Fig. 13 and 14). This suggests that the *Tc-foxQ2II* cells contribute to fast neural circuits in the central nervous system (CNS). We found these neurotransmitters expressed in the MB and CX as well (Supplementary Fig. 15 A and B).

**Table 1.**
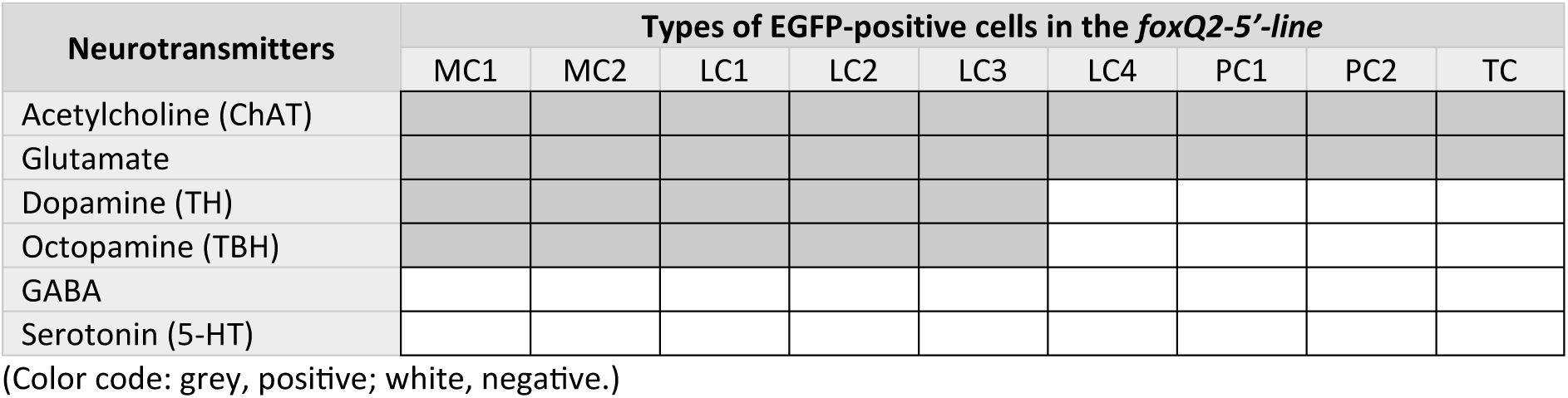
Neurotransmitter expression in different EGFP-positive cell types in the *foxQ2-5’-line* brain. Grey indicates clusters that contained at least one neuron expressing the respective transmitter. Hence, it does not mean that all cells of a cluster have a respective transmitter nor that transmitters are co-expressed in the neurons.

Five clusters included cells positive for the biogenic amines dopamine (detected by expression of TH, Tyrosine Hydroxylase) and octopamine (detected by TBH, Tyramin-β-Hydroxylase) (Table 1; Fig. 9 and 10, and Supplementary Fig. 16). Dopamine was found in projections within the MB and CX (Supplementary Fig. 15 C-C’) while TBH stained only the cell somata (data not shown).

**Figure 9.**
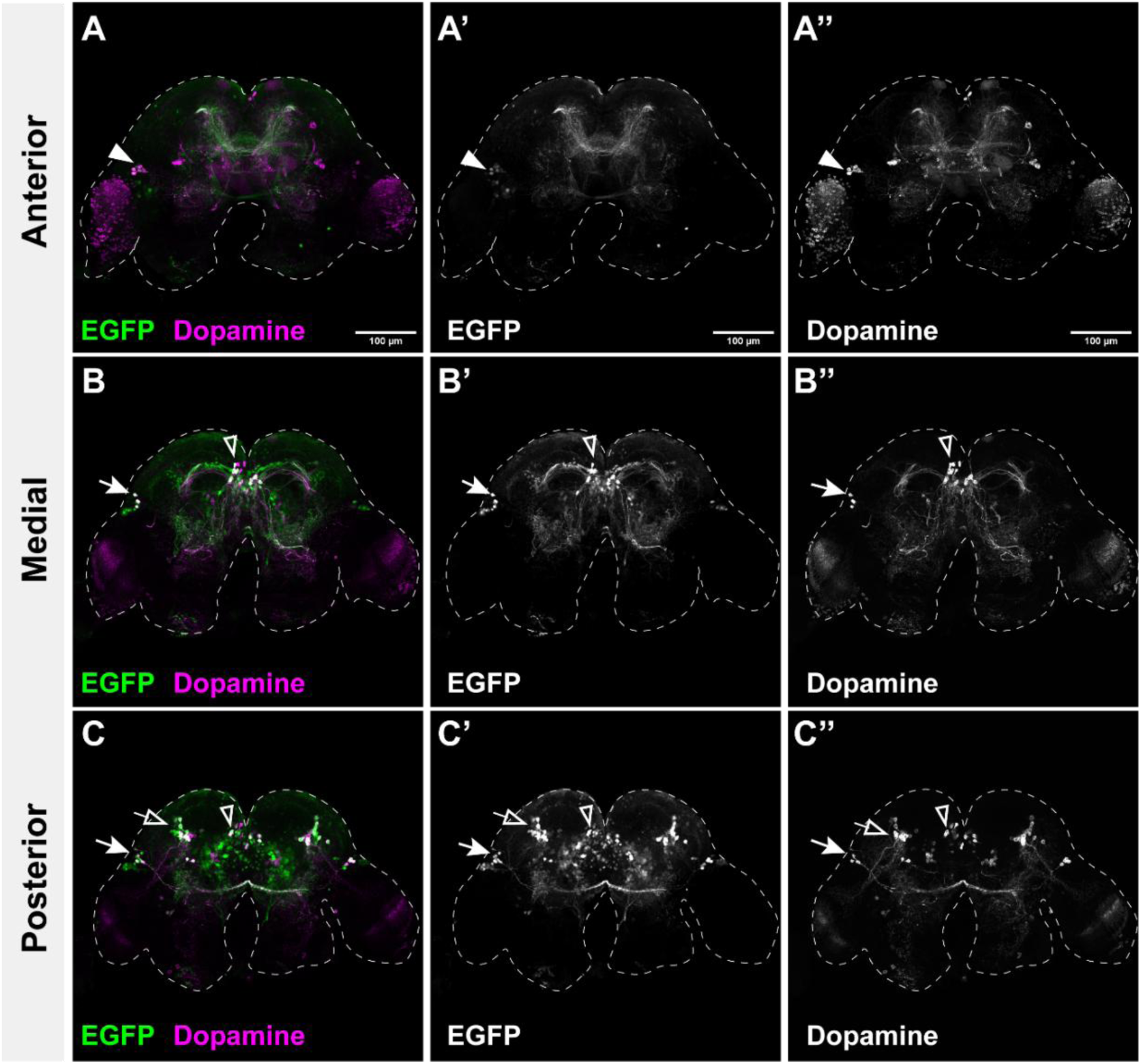
A subset of Tc-FoxQ2*-*positive neurons are dopaminergic. Co-immunostaining for GFP (green) and the dopamine marker Tyrosine Hydroxylase (TH, magenta) identifies dopaminergic subtypes among the Tc-FoxQ2-positive neuronal populations. Dashed outlines indicate the brain contour. (A-A’’) The anterior brain region containing lateral cluster 3 (LC3, white arrowheads). (A’) GFP channel. (A’’) TH (dopamine) channel. (B-B’’) A medial section showing the median cluster (MC, open arrowheads), and lateral cluster 2 (LC2, white arrows). (C-C’’) A posterior section showing the median cluster (MC, open arrowheads), lateral cluster 1 (LC1, open arrows) and lateral cluster 2 (LC2, white arrows). Scale bar: 100 μm.

**Figure 10.**
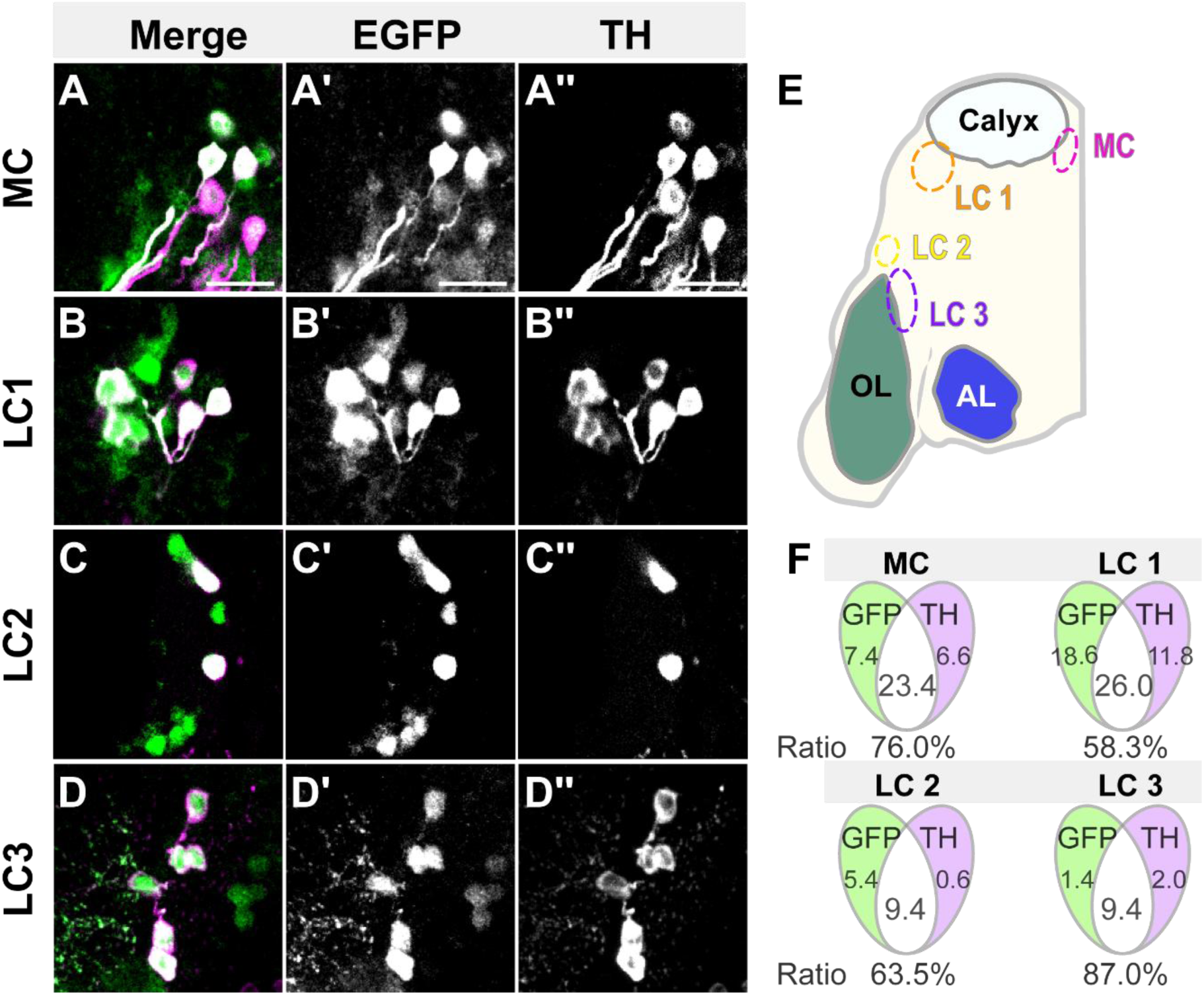
Co-expression of Tc-foxQ2II and TH (dopamine) in distinct neuronal subsets. A magnified view highlights neuronal subtypes co-expressing the dopaminergic marker Tyrosine Hydroxylase (TH) and Tc-FoxQ2/EGFP, alongside an analysis of the signal overlap ratio. (A-D’’) A representative imaging plane showing co-immunostaining for neurons expressing both EGFP (green) and TH (magenta). (E) Schematic of the left-brain hemisphere. Four Tc-FoxQ2-positive neuronal subtypes are outlined with differently colored dashed circles. (F) Venn diagrams quantify the overlap between TH-positive and Tc-FoxQ2-positive neurons for each of the four subtypes (n = 5). Scale bar: 20 μm.

Biogenic amines are involved in learning and memory and setting arousal threshholds (Davis, 2023). *This relates to the mushroom bodies’ function in olfactory memory, and the function of the central complex in visual pattern learning and goal directed navigation, respectively* (Liu et al., 2006; Wolf et al., 1998). Therefore, we focused on the analysis of dopamine in the *Tc-foxQ2II* positive neurons. First, we determined the number of cells within each cluster that were dopamine-positive and calculated the overlap ratio for each cluster (Fig. 10 A-F). In the MC (the counts for MC1 and MC2 were aggregated), overall, 76.0% of Tc-foxQ2II neurons express dopamine (Fig. 10 A-A’’ and F). In the LC1, the overlap ratio between dopamine-positive and FoxQ2-positive neurons was 58.3% (Fig. 10 B-B’’ and F) in the LC2 it was 63.5% (Fig. 10 C-C’’ and F) and in LC3 87.0% (Fig. 10 D-D’’ and F). In summary, across these clusters 60-90 % of the FoxQ2II neurons were dopamine-positive. On the other hand, we sought to determine the proportion of dopaminergic neurons that express FoxQ2. By quantifying the total number of dopamine-positive cells in the brain (excluding the optic lobe) relative to those that are also FoxQ2-positive, we found that 25.8% of dopaminergic neurons were FoxQ2-positive (n = 5).

Octopamine-labeled neurons were found in the same clusters as dopamine. For instance, octopamine staining was observed in some cells within the median cluster (MC1 and MC2) and the lateral clusters (LC1, LC2, and LC3) (Table. 1 and Supplementary Fig. 16). The portion of cells positive for octopamine within the clusters (MC, LC1, LC2 and LC3) was 61.5%, 45.2%, 58.1%, and 62.1%, respectively (Supplementary Fig. 16 H). Hence, octopamine is expressed in fewer cells compared to dopamine.

### FoxQ2 positive dopaminergic neurons can be assigned to fly neuron types and allow confirming clusters

Because only a subset of *Tc-foxQ2II* neurons are dopaminergic, we utilized TH staining as an alternative way for sparse marking. This allowed the reconstruction of clusters with greater clarity (Fig. 11 A and Supplementary Fig. 17). Analyzing the TH/EGFP double positive neurons confirmed the assignment of clusters outlined above (Compare Fig. 11 with Fig. 3 and 4). Specifically, we confirmed that the MC-cells do not cross the midline but project towards lateral regions (Fig. 11 B) and that the dopamine-positive neurons in MC1 strongly resemble PPM1 cluster dopaminergic neurons. Clusters LC2 (yellow in Fig. 11 A, E) and LC3 (green in Fig. 11 A, E) were confirmed, as well. Based on neurotransmitter content, neuronal morphology and comparison with the *Drosophila* database, we hypothesize that the dopamine-positive neurons in LC2 may correspond to PPL2c, that show similarities with the neuron with the identity CB0452 in the electron microscopy dataset (Schlegel et al., 2024), that recently has been predicted as being dopaminergic (Eckstein et al., 2024) and those in LC3 may correspond to PPL2ab (Supplementary Fig. 3) that have been identified as PPL204 in the electron microscopy dataset (Scheffer et al., 2020; Schlegel et al., 2024).

**Figure 11.**
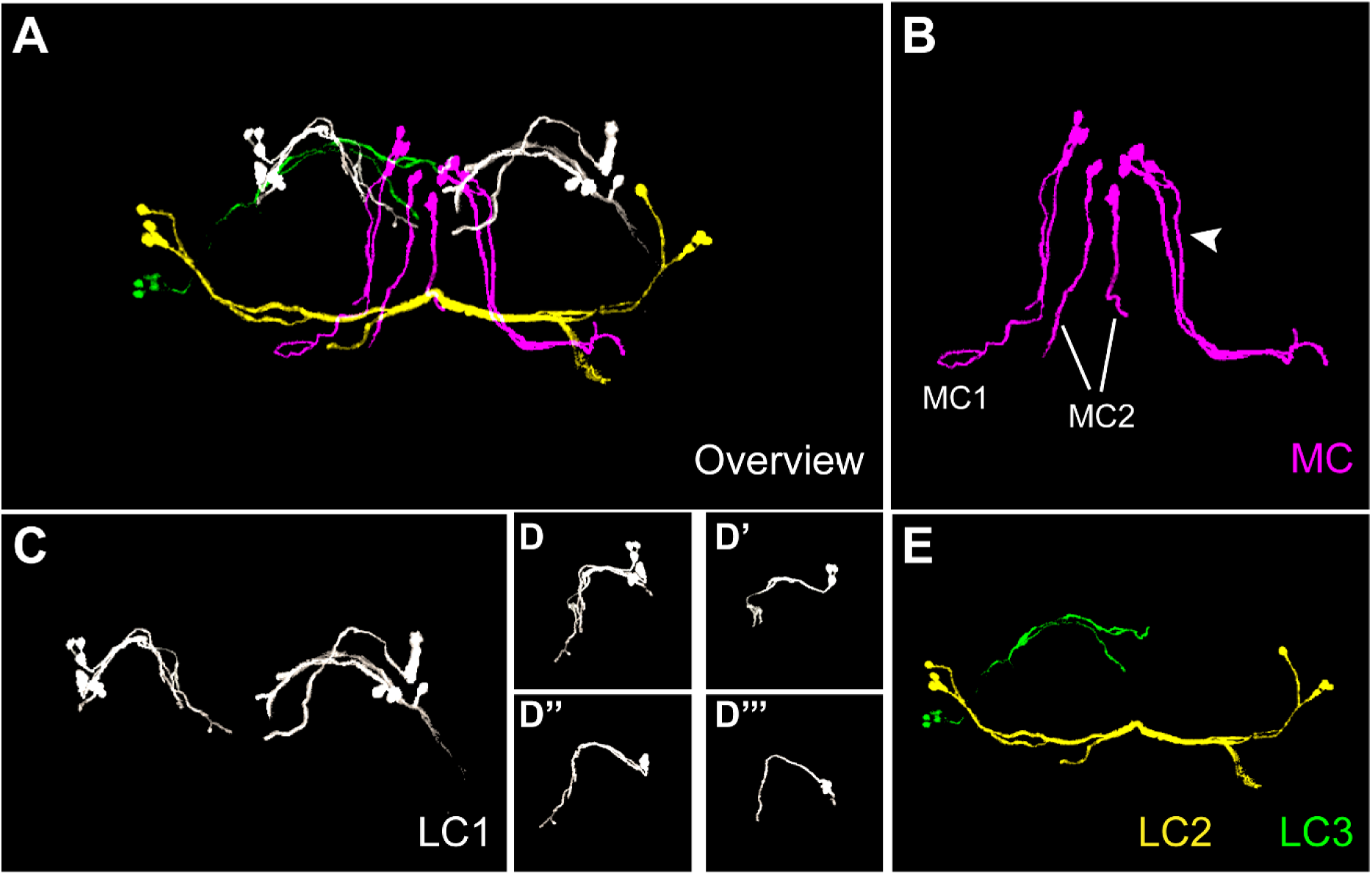
3D reconstruction of *Tc-foxQ2*-Dopaminergic neuronal projections in the brain of *foxQ2-5’-line* using VVDviewer. (A) Overview of the projections from the four neuronal subtypes. (B) Projections from the median cluster (MC). MC1 contains two long tracts (indicated by a white arrowhead), consistent with previous descriptions. (C) Projections from all subtypes within lateral cluster 1 (LC1). (D-D’’’) Individual projections of each LC1 subtype. (D) Overview of the hemi-LC1. (D’) LC1a projection (see also Fig. 5). (D’’-D’’’) LC1b projections (see also Fig. 5). (E) Projections from LC2 and LC3. For the position of the neurons within the brain see Fig. 3B/C (MC), Fig. 3E (LC2), Fig. 3D and Fig. 5 (LC1) and Fig. 4A (LC3).

With respect to the connection between MB and CX we found that all dopamine-positive cells from LC1 projected towards the CB contacting the MB on their way (Fig. 11 C-D’’’). Furthermore, we identified the previously detected LC1a and LC1b subtypes (Fig. 11 D-D’’’; compare to Fig. 5). The neurons potentially connecting CB with MB were dopamine positive and similar to the *Drosophila* PPL1 subtype.

## Discussion

### *Tc-foxQ2II* is expressed in fast-acting neurons and may be involved in the specification of subtypes of dopaminergic neurons

*foxQ2II* is an ancestral transcription factor involved in anterior patterning of most animal clades but notably was lost from vertebrates. Given its documented role in apical organ development, it is likely involved in specifying sensory and neuroendocrine cells. Indeed, other paralogs of that family have been shown to be involved in sensory cells of the cnidarian apical organ and the fish retina (Gattoni et al., 2025b; Ogawa et al., 2021). However, the contribution of *foxQ2II* marked neurons to the adult brain has remained unstudied. Therefore, our results give a first impression of *foxQ2II* function in protostome brains and provides a basis for comparison with deuterostomes. We found a wide range of different neuron types to be marked by *Tc-foxQ2II*. Nine clusters of neurons were distinguished by cell body location and projection pattern. Cell body locations ranged from median-anterior to posterior-lateral, arborizations innervated different neuropils and both ipsilateral and contralateral projections were found. Hence, neurons marked by this transcription factor have not much morphological similarity. Intriguingly, a pattern was found with respect to the neurotransmitters: Adult *Tc-foxQ2II* positive neurons seem to be predominantly fast acting as most of them express Glutamate and/or the marker for cholinergic neurons (ChAT). None of the neurons was positive for serotonin (5-HT) or GABA. The former function corresponds to studies of foxQ2 in sea urchin, where serotonergic neurons develop in a region that has lost foxQ2 expression (Yaguchi et al., 2016). Interestingly, only a subset of dopaminergic neurons was *Tc-foxQ2II* positive in *Tribolium*. We therefore hypothesize that it may be involved in the specification of dopaminergic neuron subtypes. The fact that not all dopaminergic neurons were *Tc-foxQ2II* positive and that only parts of the *Tc-foxQ2II* marked cells of any given cluster was dopamine positive indicates additional regulation, e.g. by other transcription factors and/or the Delta Notch pathway, which defines two divergent hemilineages from one neural lineage (Truman et al., 2010). Taken together, *Tc-foxQ2II* is likely part of the combinatorial specification system of adult neurons, where transcription factor “cocktails” determine cell identity. This paradigm of neural specification has been established in the insect ventral nerve cord (Skeath and Thor, 2003; Technau et al., 2006) and is thought to act in the brain as well, albeit involving different genes (Epiney et al., 2025; Özel et al., 2022; Urbach et al., 2003). To disentangle the precise regulatory function, it will be required to mark individual or small subsets of *Tc-foxQ2II* positive neurons before performing functional tests with candidate regulators.

### Stage specific expression of *Tc-foxQ2II* in columnar and tangential neurons of the central complex

The central complex (CX) is a highly complex neural structure composed of very distinct types of neurons. Among them is the class of columnar neurons, which have their cell bodies in the *pars intercerebralis* and interconnect slices of different CX subdivisions in a complex but largely conserved projection pattern. Tangential neurons in contrast connect other brain regions to certain CX subdivisions (Pfeiffer and Homberg, 2014). It came as a surprise that *Tc-foxQ2II* seems to be active in lineages giving rise to both, columnar and tangential neurons. Specifically, the *foxQ2-5’* enhancer trap activity indicated ongoing expression in the adult brain of *Tc-foxQ2II* in tangential (but not columnar) neurons of the CX. In contrast, the permanent marking by the brainbow system revealed columnar neurons as well (Supplementary Fig. 9). This indicates that the enhancer trap line had been active in neural lineages that give rise to columnar neurons at early stages but that *Tc-foxQ2II* expression was shut down in the adult brain. In line with this hypothesis, we have detected embryonic *Tc-foxQ2II* expression in putative type II neuroblasts, which are known to contribute to columnar neurons (He et al., 2019). Hence, the same transcription factor can be involved in the development of fundamentally different neuron types of one brain center such as columnar and tangential neurons. Temporally separated activity in the different lineages may be required for that multifaceted role. Interestingly, the transcription factor *retinal homeobox* also marks both, columnar and tangential neurons in *Tribolium* and *Drosophila* (Bullinger, 2024). Hence, this pattern might be more widespread and reflect a developmental principle.

### FoxQ2II marks dopaminergic neurons of the higher brain centers *mushroom bodies* and *central complex*

The mushroom bodies (MBs) and the CX are two higher order brain centers with functions in learning and memory and spatial orientation, respectively (Heisenberg, 2003; Pfeiffer and Homberg, 2014). Only recently, connections between these two neuropils were found in flies based on EM connectome and transsynaptic tracing. Specifically, some mushroom body output neurons (MBONs) seem to connect to the fan-shaped body (Kandimalla et al., 2023; Li et al., 2020; Scaplen et al., 2020) and there is evidence for such connections in *Heliconus* butterflies as well (Farnworth et al., 2025). With respect to dopaminergic neurons, in the electron microscopy dataset it appears that the PPL108 neuron was found to connect the noduli of the CX with the MBs(Scheffer et al., 2020; Schlegel et al., 2024). We find that *Tc-foxQ2II* is expressed in some dopamine positive PPL1 neurons. Our reconstructions indicate that some of these neurons have arborizations in the MBs and the fan-shaped body of the CX. We cannot rule out that there are *Tc-foxQ2II* positive dopaminergic neurons from this cluster that provide direct connectivity between these brain centers. For the time being, this has to remain a hypothesis because too many neurons were marked to make clear statements. A transgenic system to mark single cells in *Tribolium* will be required to unequivocally show such neurons. Our brainbow approach allowed for sparse marking but did not reach single cell resolution. Increasing the expression level for sensitivity and reducing the efficacy of Cre-recombination are steps that would enhance the system.

## 3.1.6 Materials and methods

| Key Resources Table |  |  |  |  |
| --- | --- | --- | --- | --- |
| Reagent type (species) or resource | Designation | Source or reference | Identifiers | Additional information |
| gene | *Tc-foxQ2II* (*Tribolium castaneum*) | iBeetle-Base<br>NCBI | TC004761<br>LOC657858 | Formerly termed *Tc-foxQ2*<br>*Drosophila* ortholog: *fd102*, CG11152 |
| strain | *San Bernardino* |  | *SB* | NCBI:txid7070 |
| genetic reagent | *foxQ2-5'-line* (*T. castaneum*, *vermillion white* background) | He et al. 2019 |  | Marks *Tc-foxQ2* positive cells with EGFP and Cre; available from the Bucher lab |
| **antibody** | *Tc-foxQ2*<br>antibody<br>(guinea pig) | He et al. 2019 |  | 1:1000 |
| **antibody** | chicken anti-GFP | Abcam | RRID: ab13970 | 1:2000 |
| **antibody** | rabbit anti-GFP | Thermo Fisher Scientific | RRID: A-11122 | 1:1000 |
| **antibody** | rabbit anti-RFP | Abcam | RRID:AB_945213 | 1:1000 |
| **antibody** | mouse anti-Synapsin | DSHB | RRID: 3C11 | 1:25 |
| **antibody** | mouse anti-Tyrosine-Hydroxylase | Immunostar | RRID: 22941 | 1:1000 |
| **antibody** | rabbit anti-Serotonin | Sigma-Aldrich, Merck KGaA | RRID: S5545 | 1:1000 |
| **antibody** | rabbit anti-GABA | Sigma-Aldrich, Merck KGaA | RRID: A2052 | 1:100 |
| **antibody** | mouse anti-Glutamate | Sigma-Aldrich | RRID: G9282 | 1:1000 |
| **antibody** | mouse anti-ChAT4B1 | DSHB | RRID: AB_528122 | 1:200 |
| **antibody** | rabbit anti-TβH | | Gift from Uwe Homberg | 1:30 |
| **antibody** | Secondary antibodies coupled with Alexa Fluor 488, 555 or 647 | Thermo Fisher Scientific | RRID: A-11039, RRID: A-11070, RRID: A-21435, RRID: A-21425, RRID: A-21430, RRID: A-21450, RRID: A-21235, RRID: A-21245 | 1:1000 |
| **antibody** | Goat anti-rabbit STAR RED | Abberior | RRID: STRED-1002-500UG | 1:1000 |
| **antibody** | Goat anti-mouse STAR ORANGE | Abberior | RRID: STORANGE-1001-500UG | 1:1000 |
| **software, algorithm** | VVD viewer | Janelia Open Science Software Initiative |  | Used for reconstructing projections from stacks of stained brains |

### Animals

*Tribolium castaneum* stocks were raised under standard conditions at 32 °C or 23 °C. The *foxQ2-5’-line* (He, 2019), an enhancer trap transgenic line generated by genome editing, was used for immunohistochemical staining, behavioral testing and RNAi experiments. The *EFA-Brainbow line* (generated using the strategy described in (Schomburg, 2010)) was used for sparse labeling of *Tc-foxQ2II* positive cells.

### Generation of the *EFA-Brainbow line*

The EFA-Brainbow construct was made starting from the Brainbow 1.1 M construct of (Livet et al., 2007). First, we disrupted the Kusabira orange fluorescent protein by digesting the original Brainbow plasmid with MluI, filling in with Klenow and re-ligating the cleaved ends, thus introducing a frameshift within the coding sequence of Kusabira. Then we cloned the the 4.5 kb NheI-SspI fragment containing the Brainbow cassette into the pSLfa1180fa plasmid (Horn and Wimmer, 2000), downstream of the EFA cis-regulatory sequence (Sarrazin et al., 2012), generating the EFA-Brainbow construct. An 8kb AscI fragment containing the EFA-Brainbow construct was then taken from this plasmid and cloned into the AscI site of pBac(3xP3-EGFP) and pBac(3xP3-DsRed) transgenesis vectors (Horn and Wimmer, 2000). The plasmids and corresponding sequences are available from Addgene (#256610 with 3xP3-EGFP eye marker and #256611 with the 3xP3-DsRed eye marker).

### Immunohistochemistry

Immunohistochemistry of larval and adult brains was performed according to a previously described protocol (Hunnekuhl VS et al., 2020) and https://doi.org/10.1007/978-1-4939-9732-9_12. The following primary antibodies were used: chicken anti-GFP (1:2000, Abcam plc; RRID: ab13970), rabbit anti-GFP (1:1000, Thermo Fisher Scientific Inc.; RRID: A-11122), rabbit anti-RFP (1:1000, Abcam; RRID:AB_945213), guinea pig anti-FoxQ2 (1:1000) (He, 2019), mouse anti-Synapsin (1:25, DSHB; RRID: 3C11), mouse anti-Tyrosine-Hydroxylase (1:1000, Immunostar Inc.; RRID: 22941), rabbit anti-Serotonin (1:1000, Sigma-Aldrich, Merck KGaA; RRID: S5545), rabbit anti-GABA (1:100, Sigma-Aldrich, Merck KGaA; RRID: A2052), mouse anti-Glutamate (1:1000, Sigma-Aldrich; RRID: G9282), mouse anti- ChAT4B1 (1:200, Developmental Studies Hybridoma Bank; RRID: AB_528122) and rabbit anti-TβH (1:30, a gift from Uwe Homberg). Secondary antibodies coupled with Alexa Fluor 488, Alexa Fluor 555 or Alexa Fluor 647 (Thermo Fisher Scientific, RRID: A-11039, RRID: A-11070, RRID: A-21435, RRID: A-21425, RRID: A-21430, RRID: A-21450, RRID: A-21235, RRID: A-21245) were used at 1:1000. STED imaging was performed with the following secondary antibodies: Goat anti-rabbit STAR RED (1:1000, Abberior, RRID: STRED-1002-500UG) and Goat anti-mouse STAR ORANGE (1:1000, Abberior, RRID: STORANGE-1001-500UG). DAPI (1:2000, Thermo Fisher Scientific Inc.; RRID: D1306) was used to stain nuclei.

### Cell counting

To quantify the number of cells in each cluster, we used the Cell counter plugin in FIJI to manually count somata of neurons in confocal image stacks of the brains analyzed in this study. Cells of interest were identified based on EGFP, mCherry or dopamine (TH) signals. Different clusters were distinguished according to cell position and projection patterns. Cell counts were performed on five standard brains.

### 3D reconstruction

The neuropil compartments of the adult brain were reconstructed using Amira software (Thermo Fisher Scientific), based on synapsin immunostaining and following the morphological framework described by Farnworth (Farnworth et al., 2022). Neuronal projections were reconstructed from confocal image stacks using VVD software (Lillvis et al., 2022).

### Image processing and documentation

Immunohistochemical samples were imaged using a Zeiss LSM 980 confocal laser scanning microscope to generate multichannel image stacks. Each stack, with a resolution of 1024 × 1024 pixels, contained 100-200 slices depending on the specimen. Stacks were processed and analyzed using FIJI (Schindelin et al., 2012), with brightness and contrast adjusted as needed. Final figures were assembled in Adobe Illustrator 2020 (Adobe).

### Materials availability statement

The original image stacks are available for download and further processing at Gro.data (https://doi.org/10.25625/FWBHWP) and AVI formats can be viewed on Figshare (doi: 10.6084/m9.figshare.30531422). The *foxQ2-5’-line* is kept and distributed upon request by the Bucher lab.

## Acknowledgements

We thank Jean Livet for providing the Brainbow 1.1 M construct and Uwe Homberg for kindly providing the rabbit anti-TβH antibody. STED-compatible secondary antibodies, goat anti-rabbit STAR RED and goat anti-mouse STAR ORANGE, were provided by Abberior. We thank Dawid Lbik for pointing out the potential of STED imaging and we acknowledge the technical support by Marita Buescher, Elke Küster and Claudia Hinners. We thank the Chinese Scholarship Council (grant 202104910052 to YP), Deutsche Forschungsgemeinschaft (grant BU1443/13-1 to GB). T.D.R. was supported by the AXA Research Fund “From Genome to Structure and Function” as a team member of the chair Kei Ito and as associate of the SFB 1451 “Key mechanisms of motor control in health and disease”.

## References

Bellen, H. J., Tong, C. and Tsuda, H. (2010). 100 years of Drosophila research and its impact on vertebrate neuroscience: a history lesson for the future. Nat. Rev. Neurosci. 11, 514–522.

Bello, B. C., Izergina, N., Caussinus, E. and Reichert, H. (2008). Amplification of neural stem cell proliferation by intermediate progenitor cells in Drosophila brain development. Neural Develop. 3, 5.

Boone, J. Q. and Doe, C. Q. (2008). Identification of Drosophila type II neuroblast lineages containing transit amplifying ganglion mother cells. Dev. Neurobiol. 68, 1185–1195.

Boyan, G., Williams, L., Legl, A. and Herbert, Z. (2010). Proliferative cell types in embryonic lineages of the central complex of the grasshopper Schistocerca gregaria. Cell Tissue Res. 341, 259–277.

Buchholz, F., Leonie Ringrose, Pierre-Olivier Angrand, Fabio Rossi, and A. Francis Stewart (1996). Different thermostabilities of FLP and Cre recombinases: implications for applied site-specific recombination. Nucleic Acids Res. 24, 4256–4262.

Bullinger, G. C. (2024). Conservation and divergence of the retinal homeobox genetic neural lineage between Drosophila melanogaster and Tribolium castaneum.

Busengdal, H. and Rentzsch, F. (2017). Unipotent progenitors contribute to the generation of sensory cell types in the nervous system of the cnidarian Nematostella vectensis. Dev. Biol. 431, 59–68.

Davis, R. L. (2023). Learning and memory using Drosophila melanogaster: a focus on advances made in the fifth decade of research. Genetics 224, iyad085.

Eckstein, N., Bates, A. S., Champion, A., Du, M., Yin, Y., Schlegel, P., Lu, A. K.-Y., Rymer, T., Finley-May, S., Paterson, T., et al. (2024). Neurotransmitter classification from electron microscopy images at synaptic sites in Drosophila melanogaster. Cell 187, 2574–2594.e23.

Epiney, D., Chaya, G. N. M., Dillon, N. R., Lai, S.-L. and Doe, C. Q. (2025). Transcriptional complexity in the insect central complex: single nuclei RNA-sequencing of adult brain neurons derived from type 2 neuroblasts. 2023.12.10.571022.

Farnworth, M. S., Eckermann, K. N. and Bucher, G. (2020). Sequence heterochrony led to a gain of functionality in an immature stage of the central complex: A fly–beetle insight. PLOS Biol. 18, e3000881.

Farnworth, M. S., Bucher, G. and Hartenstein, V. (2022). An atlas of the developing *Tribolium castaneum* brain reveals conservation in anatomy and divergence in timing to *Drosophila melanogaster*. J. Comp. Neurol. 530, 2335–2371.

Farnworth, M. S., Toh, Y. P., Loupasaki, T., Hodge, E. A., Jundi, B. el and Montgomery, S. H. (2025). Distinct evolutionary trajectories of two integration centres, the central complex and mushroom bodies, across Heliconiini butterflies. eLife 14,.

Gattoni, G., Keitley, D., Sawle, A. and Benito-Gutiérrez, E. (2025a). An ancient apical patterning system sets the position of the forebrain in chordates. Sci. Adv. 11, eadq4731.

Gattoni, G., Lin, C.-Y., York, J. R., Shew, C., Keitley, D., LaBonne, C., Yu, J.-K., Gillis, J. A. and Benito-Gutiérrez, E. (2025b). Evolutionary dynamics of FoxQ2 transcription factors across metazoans reveals three ancient paralogs. *Commun*. Biol. 9, 98.

Gehring, W. J. and Ikeo, K. (1999). Pax 6: mastering eye morphogenesis and eye evolution. Trends Genet. TIG 15, 371–377.

Hartenstein, V., Spindler, S., Pereanu, W. and Fung, S. (2008). The development of the Drosophila larval brain. Adv. Exp. Med. Biol. 628, 1–31.

Hartenstein, V., Cruz, L., Lovick, J. K. and Guo, M. (2017). Developmental analysis of the dopamine-containing neurons of the *Drosophila* brain. J. Comp. Neurol. 525, 363–379.

He, B. (2019). The role of Tc-foxQ2 in the central brain development in Tribolium castaneum.

He, B., Buescher, M., Farnworth, M. S., Strobl, F., Stelzer, E. H., Koniszewski, N. D., Muehlen, D. and Bucher, G. (2019). An ancestral apical brain region contributes to the central complex under the control of foxQ2 in the beetle Tribolium. eLife 8, e49065.

Heisenberg, M. (2003). Mushroom body memoir: from maps to models. Nat. Rev. Neurosci. 4, 266–275.

Hirth, F., Kammermeier, L., Frei, E., Walldorf, U., Noll, M. and Reichert, H. (2003). An urbilaterian origin of the tripartite brain: developmental genetic insights from Drosophila. Development 130, 2365–73.

Horn, C. and Wimmer, E. A. (2000). A versatile vector set for animal transgenesis. Dev. Genes Evol. 210, 630–637.

Hunnekuhl VS, Siemanowski J, Farnworth MS, He B, and Bucher G (2020). Immunohistochemistry and Fluorescent Whole Mount RNA In Situ Hybridization in Larval and Adult Brains of Tribolium. Methods Mol Biol 2047, 233–251.

Kandimalla, P., Omoto, J. J., Hong, E. J. and Hartenstein, V. (2023). Lineages to circuits: the developmental and evolutionary architecture of information channels into the central complex. J. Comp. Physiol. A 209, 679–720.

Kitzmann, P., Weißkopf, M., Schacht, M. I. and Bucher, G. (2017). A key role for *foxQ2* in anterior head and central brain patterning in insects. Development 144, 2969–2981.

Klingler, M. and Bucher, G. (2022). The red flour beetle T. castaneum: elaborate genetic toolkit and unbiased large scale RNAi screening to study insect biology and evolution. EvoDevo 13, 14.

Koniszewski, N. D. B., Kollmann, M., Bigham, M., Farnworth, M., He, B., Büscher, M., Hütteroth, W., Binzer, M., Schachtner, J. and Bucher, G. (2016). The insect central complex as model for heterochronic brain development—background, concepts, and tools. Dev. Genes Evol. 226, 209–219.

Kunz, T., Kraft, K. F., Technau, G. M. and Urbach, R. (2012). Origin of *Drosophila* mushroom body neuroblasts and generation of divergent embryonic lineages. Development 139, 2510–2522.

Leclère, L., Bause, M., Sinigaglia, C., Steger, J. and Rentzsch, F. (2016). Development of the aboral domain in Nematostella requires β-catenin and the opposing activities of Six3/6 and Frizzled5/8. Dev. Camb. Engl. 143, 1766–1777.

Lee, H. and Frasch, M. (2004). Survey of forkhead domain encoding genes in the *Drosophila* genome: Classification and embryonic expression patterns. Dev. Dyn. 229, 357–366.

Li, F., Lindsey, J. W., Marin, E. C., Otto, N., Dreher, M., Dempsey, G., Stark, I., Bates, A. S., Pleijzier, M. W., Schlegel, P., et al. (2020). The connectome of the adult Drosophila mushroom body provides insights into function. eLife 9, e62576.

Lillvis, J. L., Otsuna, H., Ding, X., Pisarev, I., Kawase, T., Colonell, J., Rokicki, K., Goina, C., Gao, R., Hu, A., et al. (2022). Rapid reconstruction of neural circuits using tissue expansion and light sheet microscopy. eLife 11, e81248.

Liu, G., Seiler, H., Wen, A., Zars, T., Ito, K., Wolf, R., Heisenberg, M. and Liu, L. (2006). Distinct memory traces for two visual features in the Drosophila brain. Nature 439, 551–556.

Livet, J., Weissman, T. A., Kang, H., Draft, R. W., Lu, J., Bennis, R. A., Sanes, J. R. and Lichtman, J. W. (2007). Transgenic strategies for combinatorial expression of fluorescent proteins in the nervous system. Nature 450, 56–62.

Mao, Z. and Davis, R. L. (2009). Eight different types of dopaminergic neurons innervate the Drosophila mushroom body neuropil: anatomical and physiological heterogeneity. Front. Neural Circuits 3,.

Marlow, H., Tosches, M. A., Tomer, R., Steinmetz, P. R., Lauri, A., Larsson, T. and Arendt, D. (2014). Larval body patterning and apical organs are conserved in animal evolution. BMC Biol. 12, 7.

Miyares, R. L. and Lee, T. (2019). Temporal control of Drosophila central nervous system development. Curr. Opin. Neurobiol. 56, 24–32.

Namiki, S., Dickinson, M. H., Wong, A. M., Korff, W. and Card, G. M. (2018). The functional organization of descending sensory-motor pathways in Drosophila. eLife 7, e34272.

Ogawa, Y., Shiraki, T., Fukada, Y. and Kojima, D. (2021). Foxq2 determines blue cone identity in zebrafish. Sci. Adv. 7, eabi9784.

Özel, M. N., Gibbs, C. S., Holguera, I., Soliman, M., Bonneau, R. and Desplan, C. (2022). Coordinated control of neuronal differentiation and wiring by sustained transcription factors. Science 378, eadd1884.

Pfeiffer, K. and Homberg, U. (2014). Organization and Functional Roles of the Central Complex in the Insect Brain. Annu. Rev. Entomol. 59, 165–184.

Posnien, N., Schinko, J. B., Kittelmann, S. and Bucher, G. (2010). Genetics, development and composition of the insect head – A beetle’s view. Arthropod Struct. Dev. 39, 399–410.

Posnien, N., Koniszewski, N. and Bucher, G. (2011). Insect Tc-six4 marks a unit with similarity to vertebrate placodes. Dev. Biol. 350, 208–216.

Posnien, N., Hunnekuhl, V. S. and Bucher, G. (2023). Gene expression mapping of the neuroectoderm across phyla – conservation and divergence of early brain anlagen between insects and vertebrates. eLife 12, e92242.

Rethemeier, S., Fritzsche, S., Mühlen, D., Bucher, G. and Hunnekuhl, V. S. (2025). Differences in size and number of embryonic type II neuroblast lineages correlate with divergent timing of central complex development between beetle and fly. eLife 13, RP99717.

Sarrazin, A. F., Peel, A. D. and Averof, M. (2012). A Segmentation Clock with Two-Segment Periodicity in Insects. Science.

Scaplen, K. M., Talay, M., Nunez, K. M., Salamon, S., Waterman, A. G., Gang, S., Song, S. L., Barnea, G. and Kaun, K. R. (2020). Circuits that encode and guide alcohol-associated preference. eLife 9, e48730.

Schacht, M. I., Schomburg, C. and Bucher, G. (2020). six3 acts upstream of foxQ2 in labrum and neural development in the spider Parasteatoda tepidariorum. Dev. Genes Evol. 230, 95–104.

Scheffer, L. K., Xu, C. S., Januszewski, M., Lu, Z., Takemura, S.-Y., Hayworth, K. J., Huang, G. B., Shinomiya, K., Maitlin-Shepard, J., Berg, S., et al. (2020). A connectome and analysis of the adult Drosophila central brain. eLife 9, e57443.

Schindelin, J., Arganda-Carreras, I., Frise, E., Kaynig, V., Longair, M., Pietzsch, T., Preibisch, S., Rueden, C., Saalfeld, S., Schmid, B., et al. (2012). Fiji: an open-source platform for biological-image analysis. Nat. Methods 9, 676–682.

Schlegel, P., Yin, Y., Bates, A. S., Dorkenwald, S., Eichler, K., Brooks, P., Han, D. S., Gkantia, M., Dos Santos, M., Munnelly, E. J., et al. (2024). Whole-brain annotation and multi-connectome cell typing of Drosophila. Nature 634, 139–152.

Scholtz, G. and Edgecombe, G. D. (2006). The evolution of arthropod heads: reconciling morphological, developmental and palaeontological evidence. Dev Genes Evol 216, 395–415.

Schomburg, C. (2010). Etablierung des Brainbow-Systems in Tribolium.

Simeone, A., Gulisano, M., Acampora, D., Stornaiuolo, A., Rambaldi, M. and Boncinelli, E. (1992). Two vertebrate homeobox genes related to the Drosophila empty spiracles gene are expressed in the embryonic cerebral cortex. Embo J 11, 2541–50.

Sinigaglia, C., Busengdal, H., Leclère, L., Technau, U. and Rentzsch, F. (2013). The Bilaterian Head Patterning Gene six3/6 Controls Aboral Domain Development in a Cnidarian. PLoS Biol. 11, e1001488.

Sinigaglia, C., Almazán, A., Lebel, M., Sémon, M., Gillet, B., Hughes, S., Edsinger, E., Averof, M. and Paris, M. (2022). Distinct gene expression dynamics in developing and regenerating crustacean limbs. Proc. Natl. Acad. Sci. U. S. A. 119, e2119297119.

Skeath, J. B. and Thor, S. (2003). Genetic control of Drosophila nerve cord development. Curr. Opin. Neurobiol. 13, 8–15.

Technau, G. M., Berger, C. and Urbach, R. (2006). Generation of cell diversity and segmental pattern in the embryonic central nervous system of Drosophila. Dev Dyn 235, 861–9.

Tomer, R., Denes, A. S., Tessmar-Raible, K. and Arendt, D. (2010). Profiling by image registration reveals common origin of annelid mushroom bodies and vertebrate pallium. Cell 142, 800–9.

Truman, J. W. and Riddiford, L. M. (2023). Drosophila postembryonic nervous system development: a model for the endocrine control of development. Genetics 223, iyac184.

Truman, J. W., Moats, W., Altman, J., Marin, E. C. and Williams, D. W. (2010). Role of Notch signaling in establishing the hemilineages of secondary neurons in *Drosophila melanogaster*. Development 137, 53–61.

Urbach, R. and Technau, G. M. (2003a). Early steps in building the insect brain: neuroblast formation and segmental patterning in the developing brain of different insect species. Arthropod Struct. Dev. 32, 103–123.

Urbach, R. and Technau, G. M. (2003b). Segment polarity and DV patterning gene expression reveals segmental organization of the *Drosophila* brain. Development 130, 3607–3620.

Urbach, R. and Technau, G. M. (2004). Neuroblast formation and patterning during early brain development in *Drosophila*. BioEssays 26, 739–751.

Urbach, R., Schnabel, R. and Technau, G. M. (2003). The pattern of neuroblast formation, mitotic domains and proneural gene expression during early brain development in *Drosophila*. Development 130, 3589–3606.

Walsh, K. T. and Doe, C. Q. (2017). Drosophila embryonic type II neuroblasts: origin, temporal patterning, and contribution to the adult central complex. Dev. Camb. Engl. 144, 4552–4562.

Willig, K. I., Kellner, R. R., Medda, R., Hein, B., Jakobs, S. and Hell, S. W. (2006). Nanoscale resolution in GFP-based microscopy. Nat. Methods 3, 721–723.

Wolf, R., Wittig, T., Liu, L., Wustmann, G., Eyding, D. and Heisenberg, M. (1998). Drosophila Mushroom Bodies Are Dispensable for Visual, Tactile, and Motor Learning. Learn. Mem. 5, 166–178.

Yaguchi, S., Yaguchi, J., Angerer, R. C. and Angerer, L. M. (2008). A Wnt-FoxQ2-Nodal Pathway Links Primary and Secondary Axis Specification in Sea Urchin Embryos. Dev. Cell 14, 97–107.

Yaguchi, J., Takeda, N., Inaba, K. and Yaguchi, S. (2016). Cooperative Wnt-Nodal Signals Regulate the Patterning of Anterior Neuroectoderm. PLoS Genet. 12, e1006001.

