## Supplemental figures and data for "*foxQ2* marks fast-acting interneurons including dopaminergic neurons of *mushroom bodies* and *central complex* in the beetle *T. castaneum*"

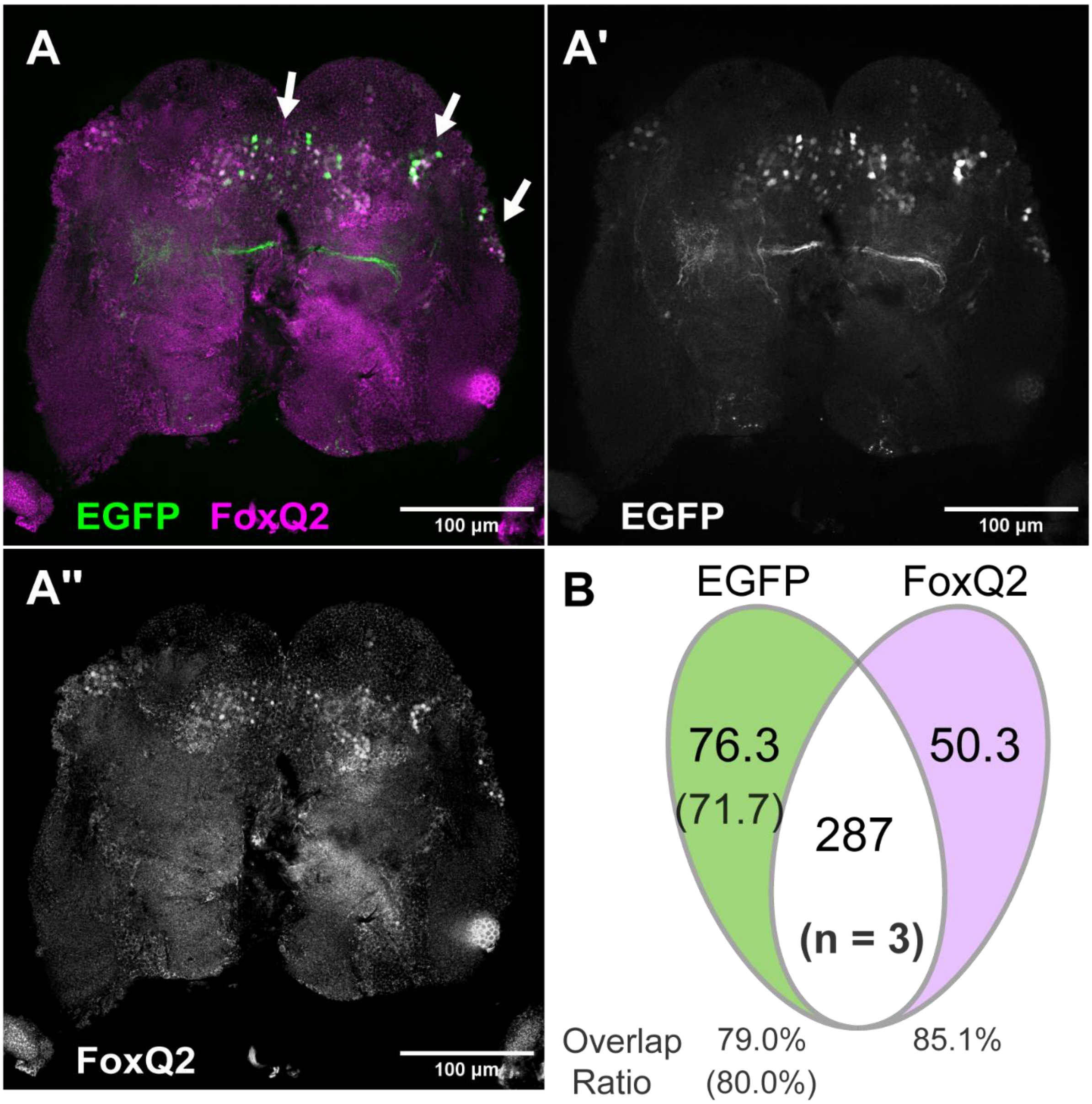


**Supplementary Figure 1. Characterization of the foxQ2-5'-imaging line** Tc-FoxQ2 showed a high degree of overlap with EGFP in adult brains in co-immunostaining of brain sections using antibodies against FoxQ2 and EGFP. (A) Cells co-expressing FoxQ2 and EGFP are indicated by white arrows. Green represents EGFP, while magenta represents FoxQ2. (B) The Venn diagram illustrates the number of cells expressing only EGFP, only FoxQ2, and those that are double-positive for EGFP and FoxQ2. The proportion of co-expressing cells relative to all EGFP- or FoxQ2-expressing cells was 79.0% for EGFP-positive cells and 85,1% for FoxQ2 positive cells. When removing the completely Tc-FoxQ2 negative AC cluster from the count (white star in Fig. 2), 80.0% of the EGFP positive cells were Tc-FoxQ2 positive), while 85.1% of FoxQ2-expressing cells overlapped with EGFP (n = 3). Scale bar: 100 µm.

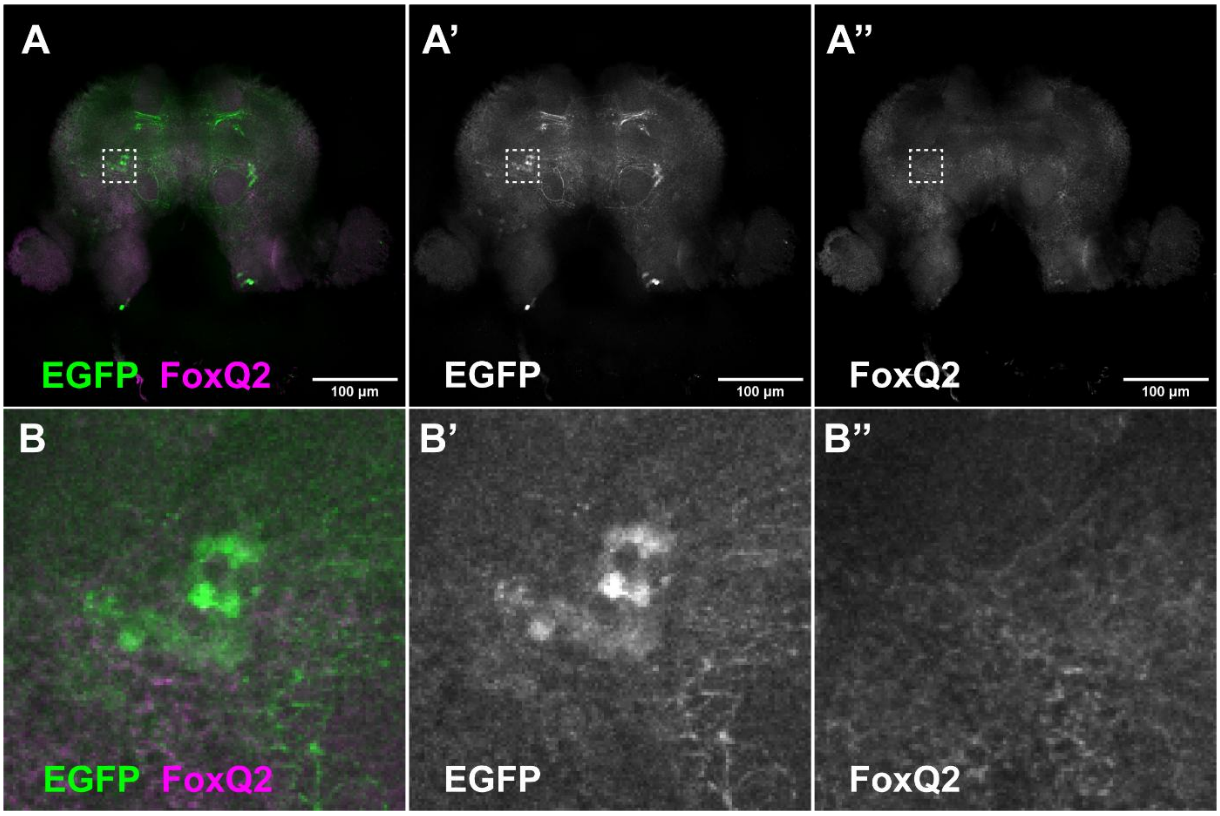


**Supplementary Figure 2. The Anterior Cluster is not marked by the FoxQ2 antibody and only expresses EGFP** (A-A'') Overview of the anterior section of the adult brain stained with EGFP and FoxQ2 antibodies. EGFP is shown in green, and FoxQ2 in magenta. (B-B'') A magnified view of the white-dashed square. The anterior cluster expresses only EGFP. Therefore, we did not consider these cells further. Scale bar: 100 µm.

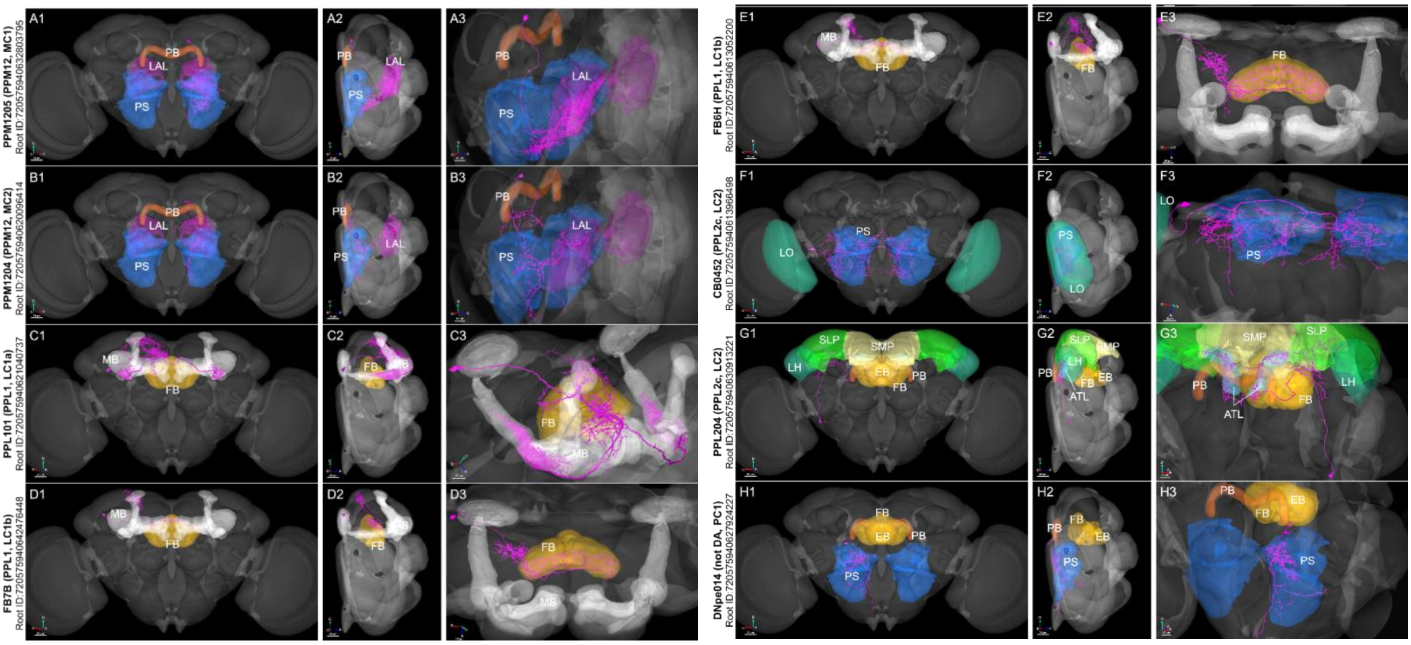


**Supplementary Figure 3. Suggested homologous neurons in *Drosophila melanogaster*.** (A-A’) Dopaminergic neuron PPM1. (B-B’) Glutamatergic neuron LAL131a. (C-C’) Dopaminergic neuron PPL1. (D-D’) Dopaminergic neuron CB0452. (E-E’) Dopaminergic neuron PPL2. (F-F’) Acetylcholinergic neuron DNpe014. (Information from https://codex.flywire.ai/, neuropils and neurons from https://v2.virtualflybrain.org/.) (I-M) Copy of figure 11 from the main text for comparison with fly neurons. See figure legend 11 for details.

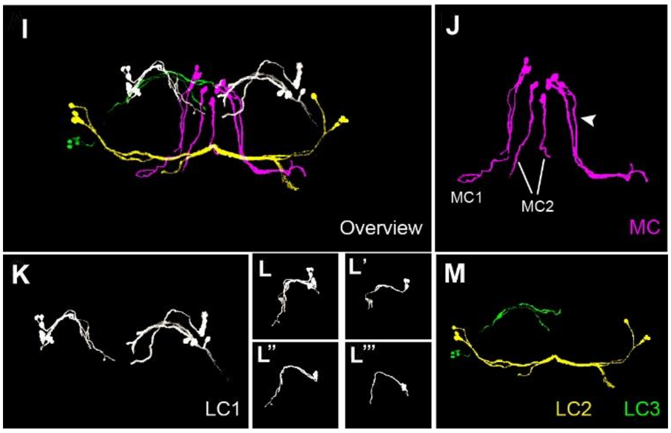

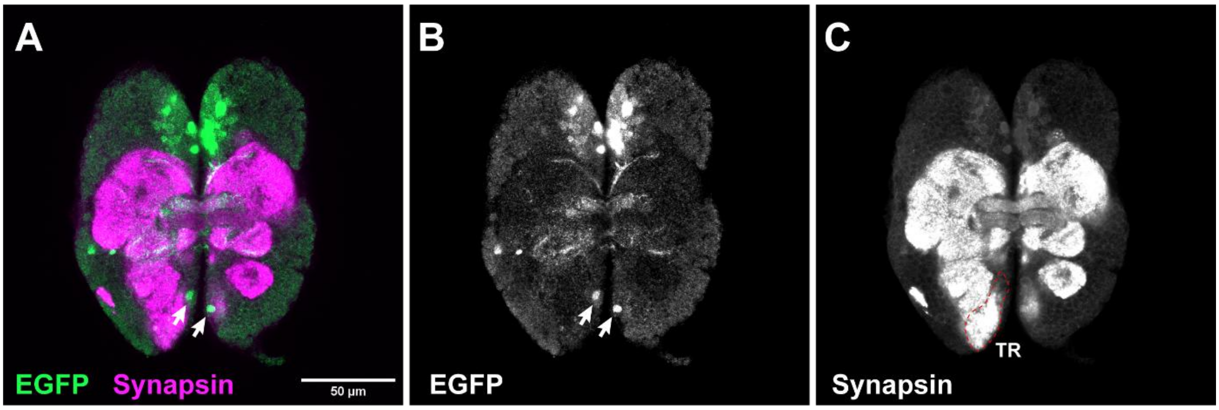


**Supplementary Figure 4.** TC emerges in the tritocerebrum (TR) of the first instar larval (L1) brain. (A) Larval brain from *foxQ2-5' line* stained with EGFP and synapsin antibodies. White arrows in (A) and (B) indicate the EGFP-positive cells located in the TR. (C) highlights the TR component (1), outlined with a red dashed line. Scale bar: 50 µm.

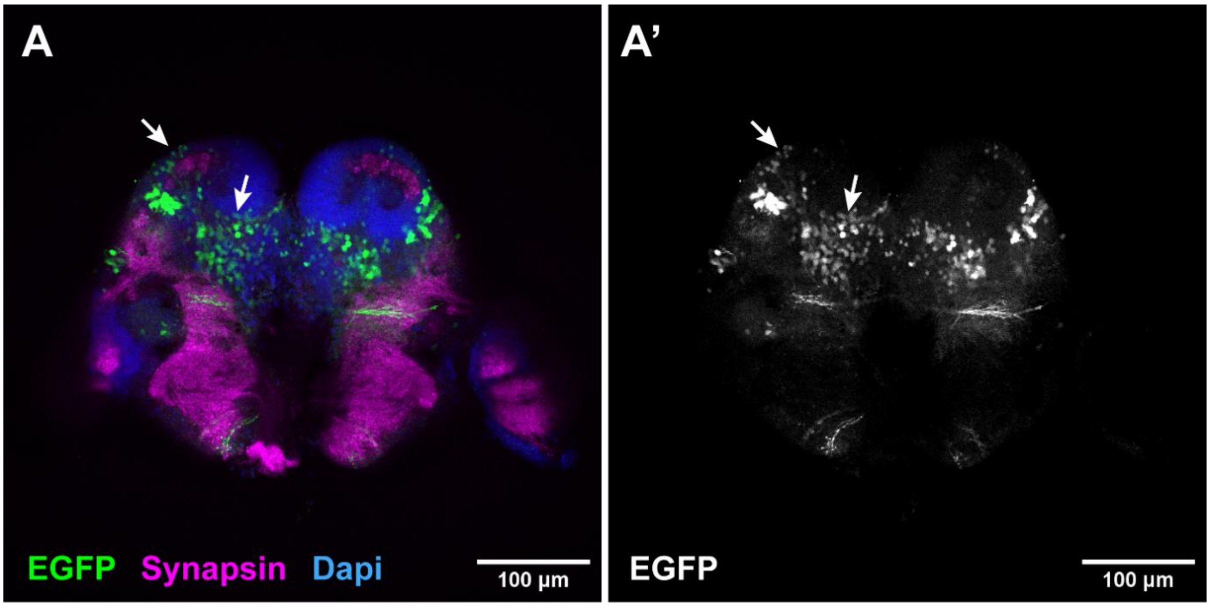


**Supplementary Figure 5. EGFP positive cells without discernible projections**. (A-A') Posterior view of the brain in the *foxQ2-5' line*. The cells indicated by white arrows lacked identifiable projections and were therefore not reconstructed. Scale bar: 100 µm.

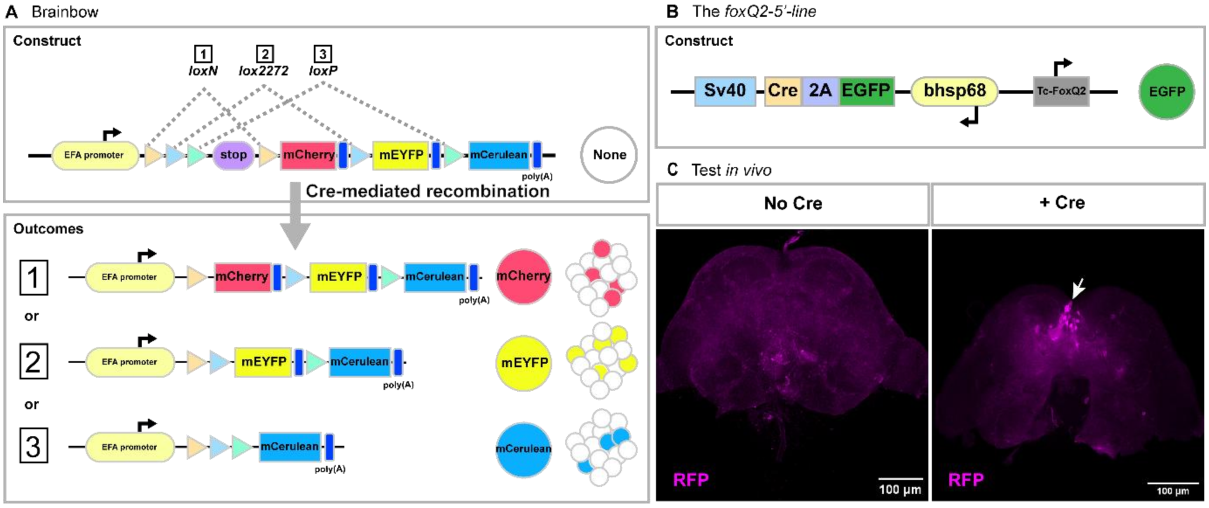


**Supplementary Figure 6.** Brainbow: stochastic recombination using incompatible lox variants. (A) In the Brainbow system, Cre randomly selects different excision events (1 - 3) based on incompatible lox sites. Before Cre activity, no fluorescence signal is detected in the brain due to the presence of a stop codon. Recombination switches expression to mCherry (1), mEYFP (2), or mCerulean (3). (B) In the *foxQ2-5' –line*, EGFP fluorescent protein and Cre recombinase are included. The enhancer activates the expression of FoxQ2, EGFP, and Cre. Since mEYFP and mCerulean fluoresce similarly to EGFP, mCherry is used as a marker to label cells in the crossed brain. (C) Before Cre activity, RFP is not detected in the Brainbow brain. When the Brainbow line is crossed with the *foxQ2-5' line*, recombination activates the expression of mCherry, resulting in RFP is detected in the brain. Scale bar: 100 µm. [EFA (3391 bp) is derived from OX466061.1.]. See Supplementary Data for specific sequence information.

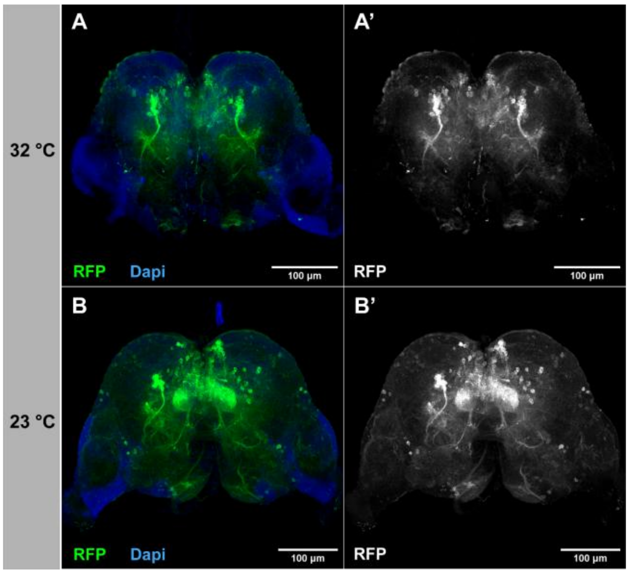


**Supplementary Figure 7.** Brains grown at 32 °C and 23 °C, stained by immunohistochemistry (IHC). (A-A') Brain raised at 32 °C stained by IHC. (B-B') Brain raised at 23 °C stained by IHC. Scale bar: 100 µm.

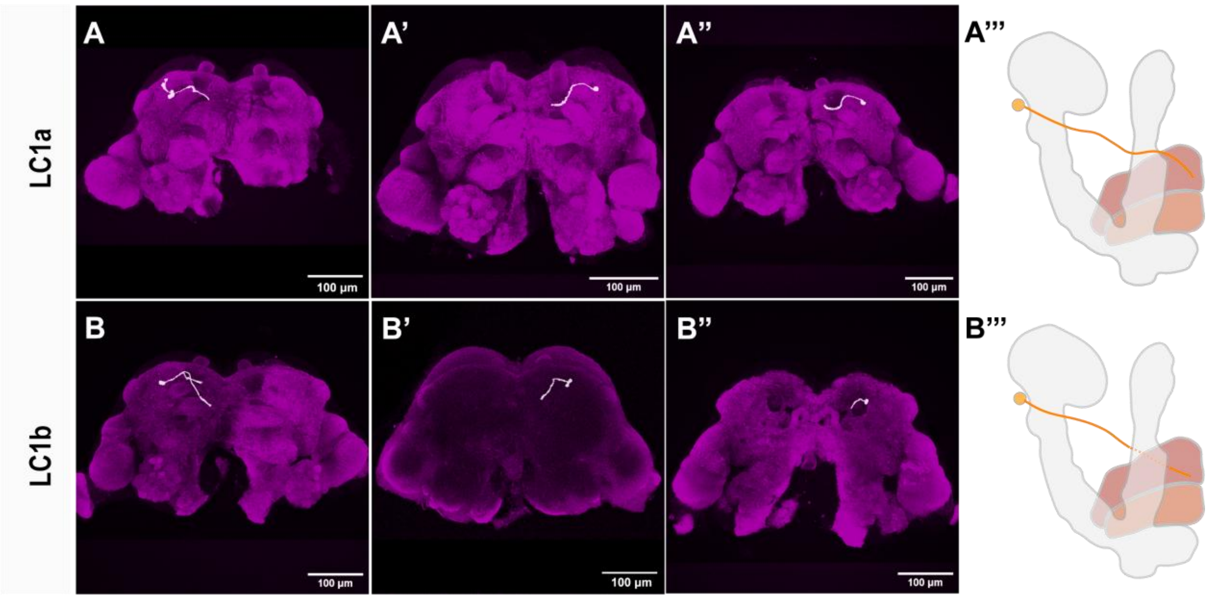


**Supplementary Figure 8.** Two LC1 subclusters are sparsely marked with the Brainbow technique. (A-A’’) LC1a axons bypass the vertical lobe of mushroom bodies and descend in three different brains. (A’’’) Schematic of LC1a projection patterns. (B-B’’) LC1b projects directly to the central body region. (B’’’) Schematic of LC1b projection patterns. Magenta represents Synapsin. (n=6). Scale bar: 100 µm.

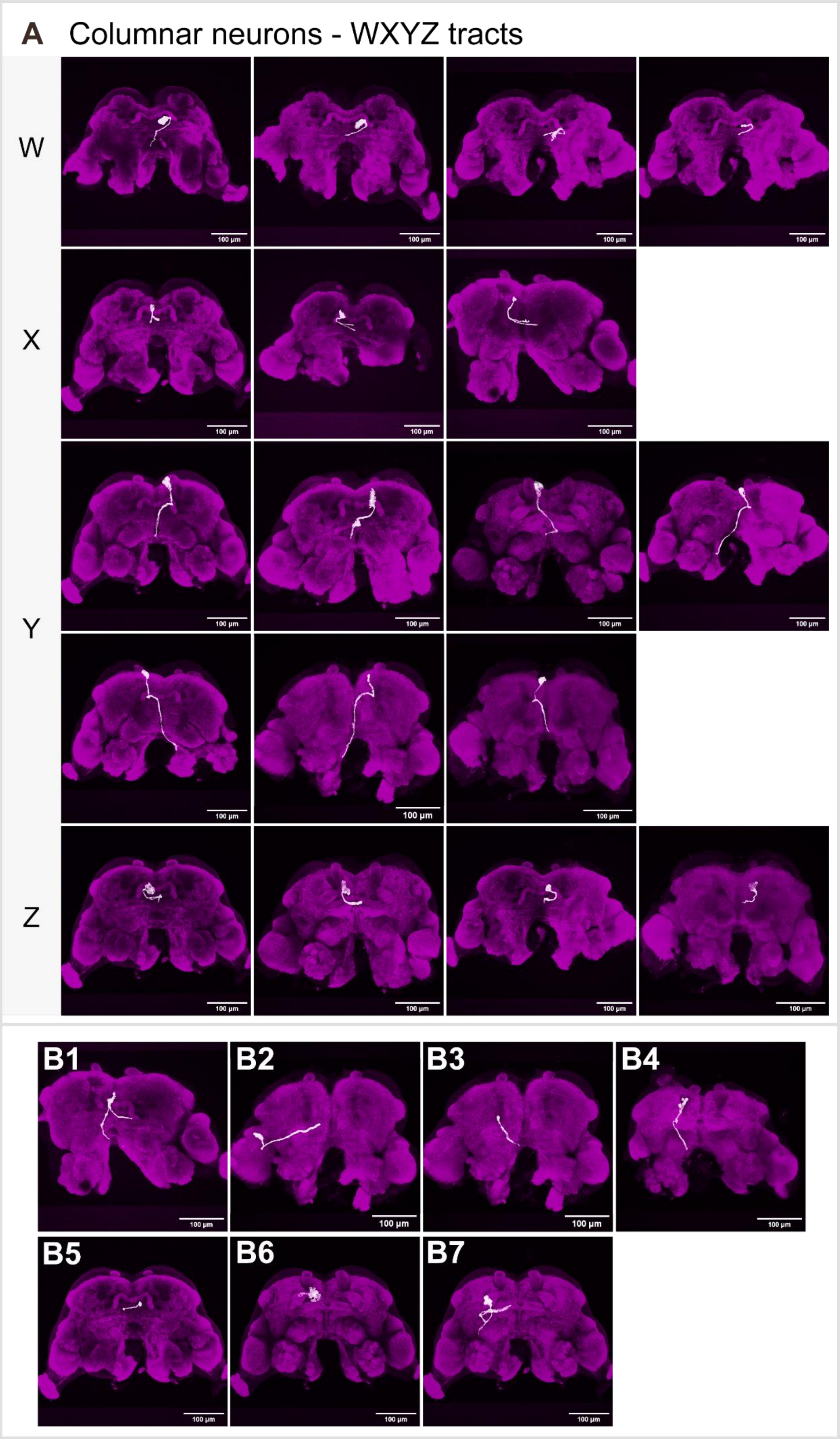


**Supplementary Figure 9. Brainbow marked cells that are not marked by adult Tc-foxQ2 expression** (A) Columnar neurons (WXYZ tracts) are sparsely labeled by Brainbow. (B) Some neurons were visualized only once in this Figure. (B1-B5) These neurons were detected only once. While (B6-B7) These neurons were consistently observed in every brain. Scale bar: 100 µm.

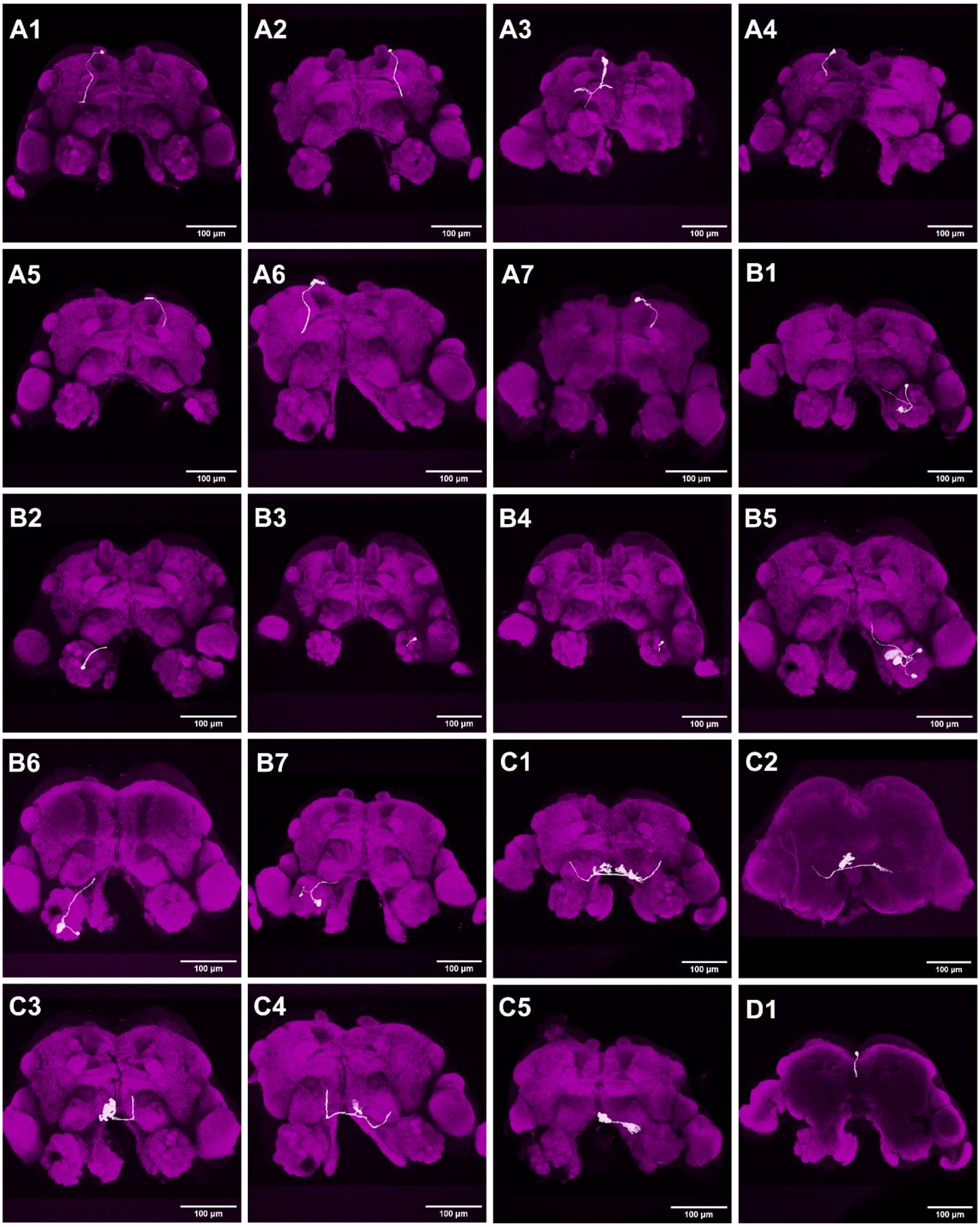


(figure continued on next page)

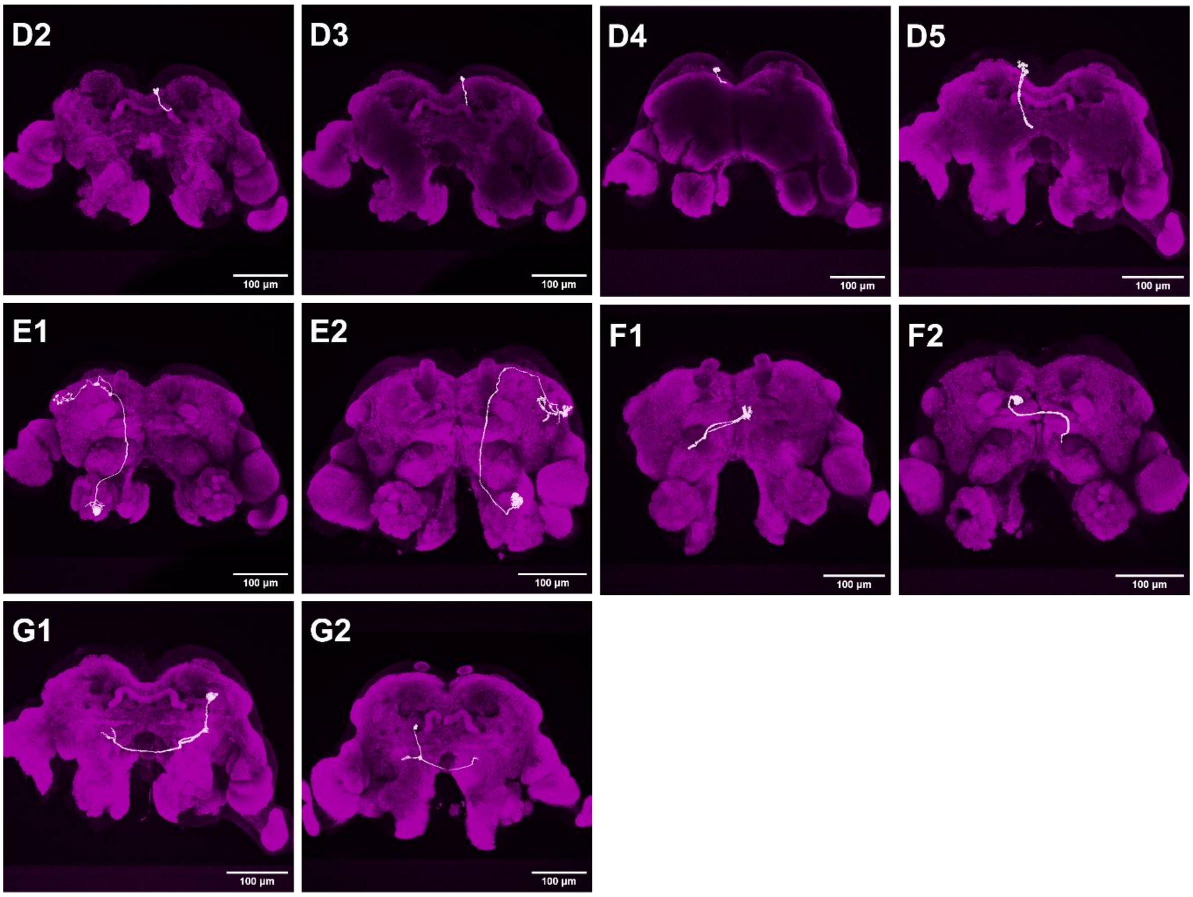


**Supplementary Figure 10. Brainbow marked cells that are not marked by adult Tc-foxQ2 expression.** Individual cells marked by sparse Brainbow technique. (A1-A7) A cell located in the posterior-dorsal region extends from the vertical lobe (VL) of the mushroom bodies to the anterior-ventral region. (B1-B7) Cells are located in and innervate the antennal lobe. (C1-C5) Cells are located in the ventral part of the brain. (D1-D5) Cells are located in the superior medial protocerebrum (SMP) and project from the dorsal to the ventral part of the brain. (E1-E2) A projection connects the lateral horn (LH) and the antennal lobe (AL). (F1-F2) Cells are located in the medial region of the brain and cross the midline. (G1-G2) Cells are located in the posterior region of the brain and cross the midline. Scale bar: 100 µm.

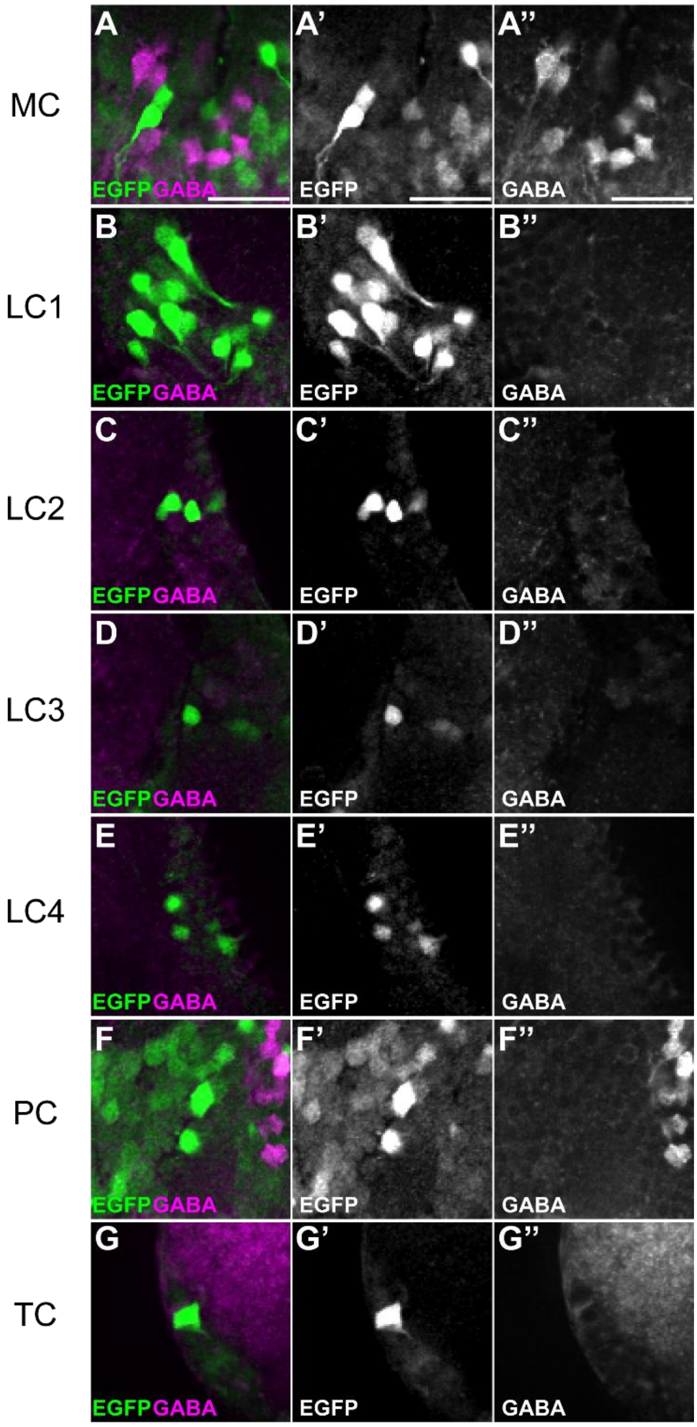


**Supplementary Figure 11.** GABA is not co-expressed with FoxQ2-positive cells. (A-A’’) In median cluster (MC), GABA and FoxQ2-positive cells are intermingled in their expression. (B-B’’) Lateral cluster 1 (LC1) shows expression of EGFP only. (C-C’’) In Lateral cluster 2 (LC2), no overlap was detected between GABA and FoxQ2 signals. (D-D’’) Lateral cluster 3 (LC3) expresses only EGFP. (E-E’’) In Lateral cluster 4 (LC4), GABA and FoxQ2 are not co-expressed in the same cells. (F-F’’) In posterior cluster (PC), GABA and FoxQ2-positive cells are intermingled in their expression. (G-G’’) The tritocerebrum cluster (TC) expresses only EGFP. Scale bar: 20 µm.

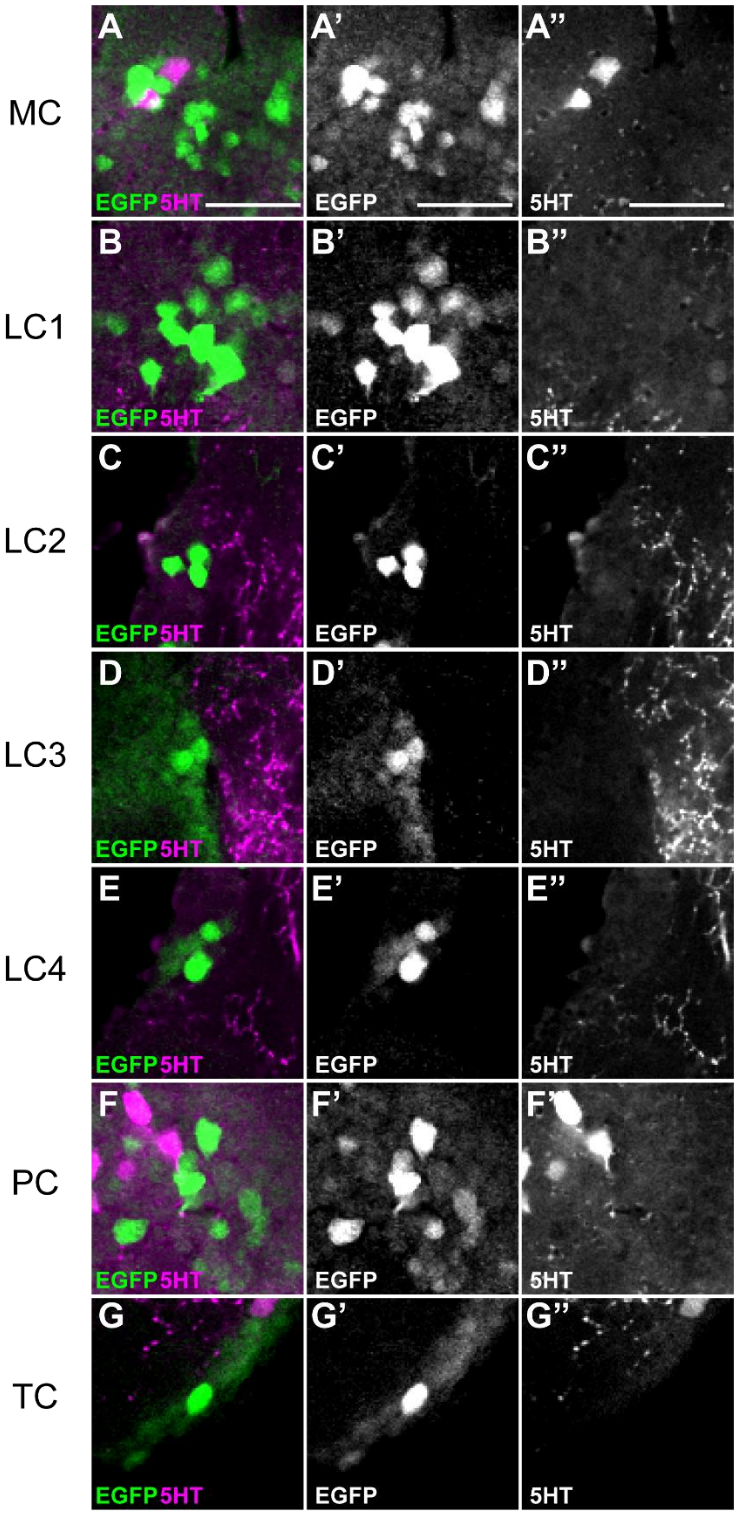


**Supplementary Figure 12.** Serotonin (5-HT) is not co-expressed with FoxQ2-positive cells. (A-A'') In median cluster (MC), 5-HT and FoxQ2-positive cells are intermingled in their expression. (B-B'') Lateral cluster 1 (LC1) shows expression of EGFP only. (C-C'') In Lateral cluster 2 (LC2), no overlap was detected between 5-HT and FoxQ2 signals. (D-D'') Lateral cluster 3 (LC3) expresses only EGFP. (E-E'') In Lateral cluster 4 (LC4), 5-HT and FoxQ2 are not co-expressed in the same cells. (F-F'') In posterior cluster (PC), 5-HT and FoxQ2-positive cells are intermingled in their expression. (G-G'') The tritocerebrum cluster (TC) expresses only EGFP. Scale bar: 20 µm.

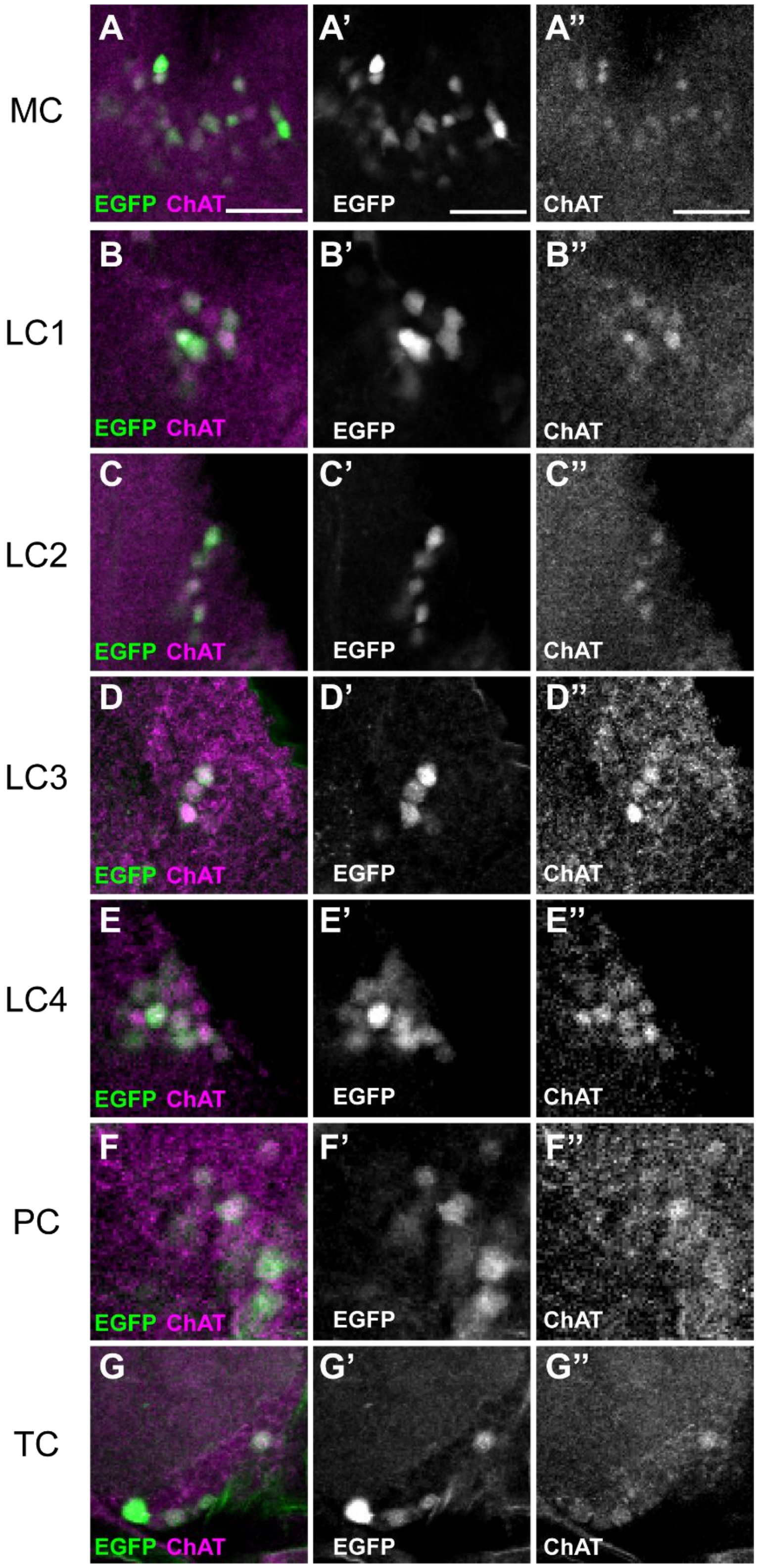


**Supplementary Figure 13.** Choline Acetyltransferase (ChAT) is expressed in all FoxQ2-positive subclusters. (A-A'') The median cluster (MC), including MC1 and MC2, expresses both ChAT and EGFP. (B-B'') Lateral cluster 1 (LC1) expresses both ChAT and EGFP. (C-C'') In Lateral cluster 2 (LC2), ChAT and EGFP largely overlap. (D-D'') Lateral cluster 3 (LC3) expresses both ChAT and EGFP. (E-E'') In Lateral cluster 4 (LC4), ChAT and EGFP largely overlap. (F-F'') The posterior cluster (PC) expresses both ChAT and EGFP. (G-G'') The tritocerebrum cluster (TC) expresses both ChAT and EGFP. Scale bar: 20 µm.

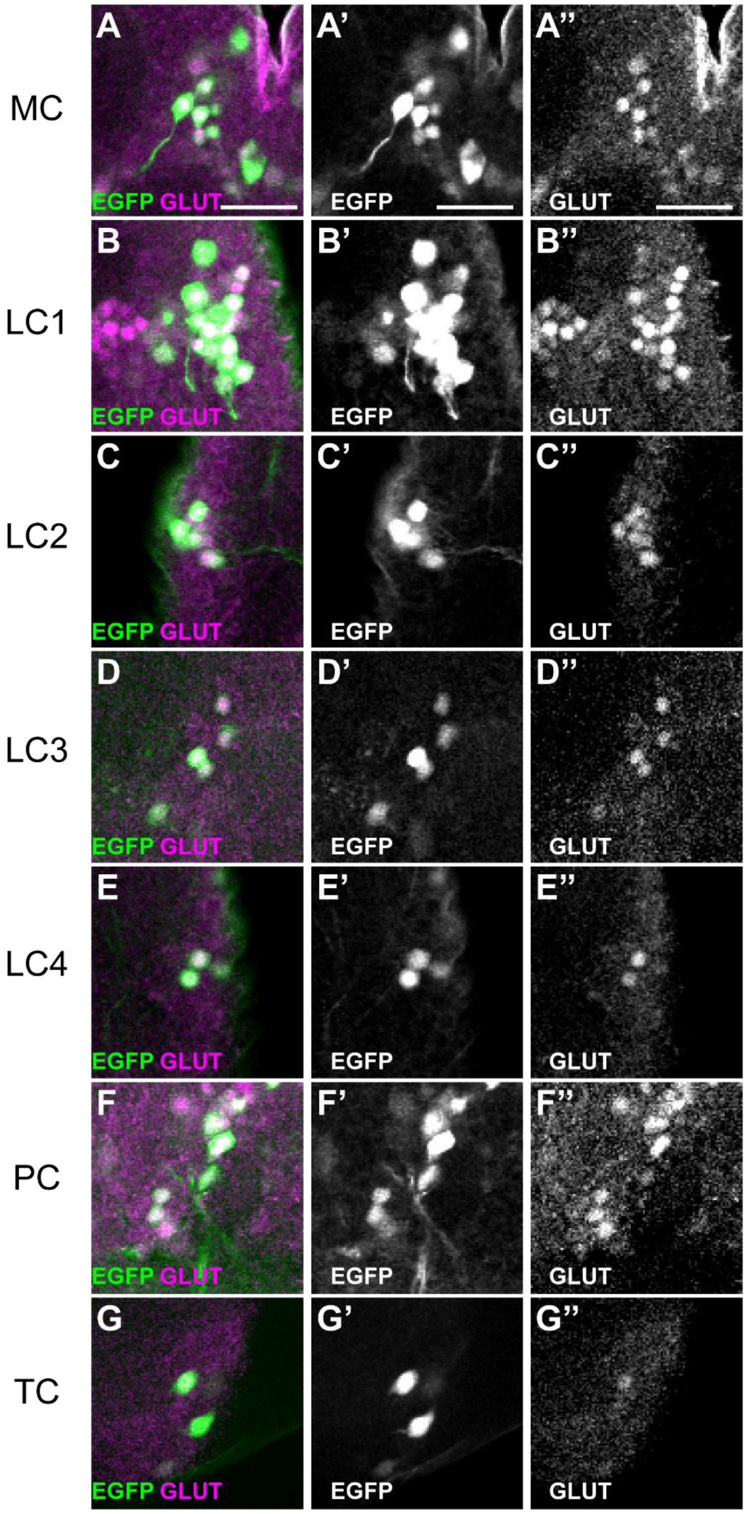


**Supplementary Figure 14.** Glutamate (GLUT) is expressed in all FoxQ2-positive subclusters. (A-A'') The median cluster (MC), including MC1 and MC2, expresses both GLUT and EGFP. (B-B'') In Lateral cluster 1 (LC1), GLUT and EGFP largely overlap. (C-C'') In Lateral cluster 2 (LC2), GLUT and EGFP largely overlap. (D-D'') Lateral cluster 3 (LC3) expresses both GLUT and EGFP. (E-E'') Lateral cluster 4 (LC4) expresses both GLUT and EGFP. (F-F'') The posterior cluster (PC) expresses both GLUT and EGFP. (G-G'') The tritocerebrum cluster (TC) expresses both GLUT and EGFP. Scale bar: 20 µm.

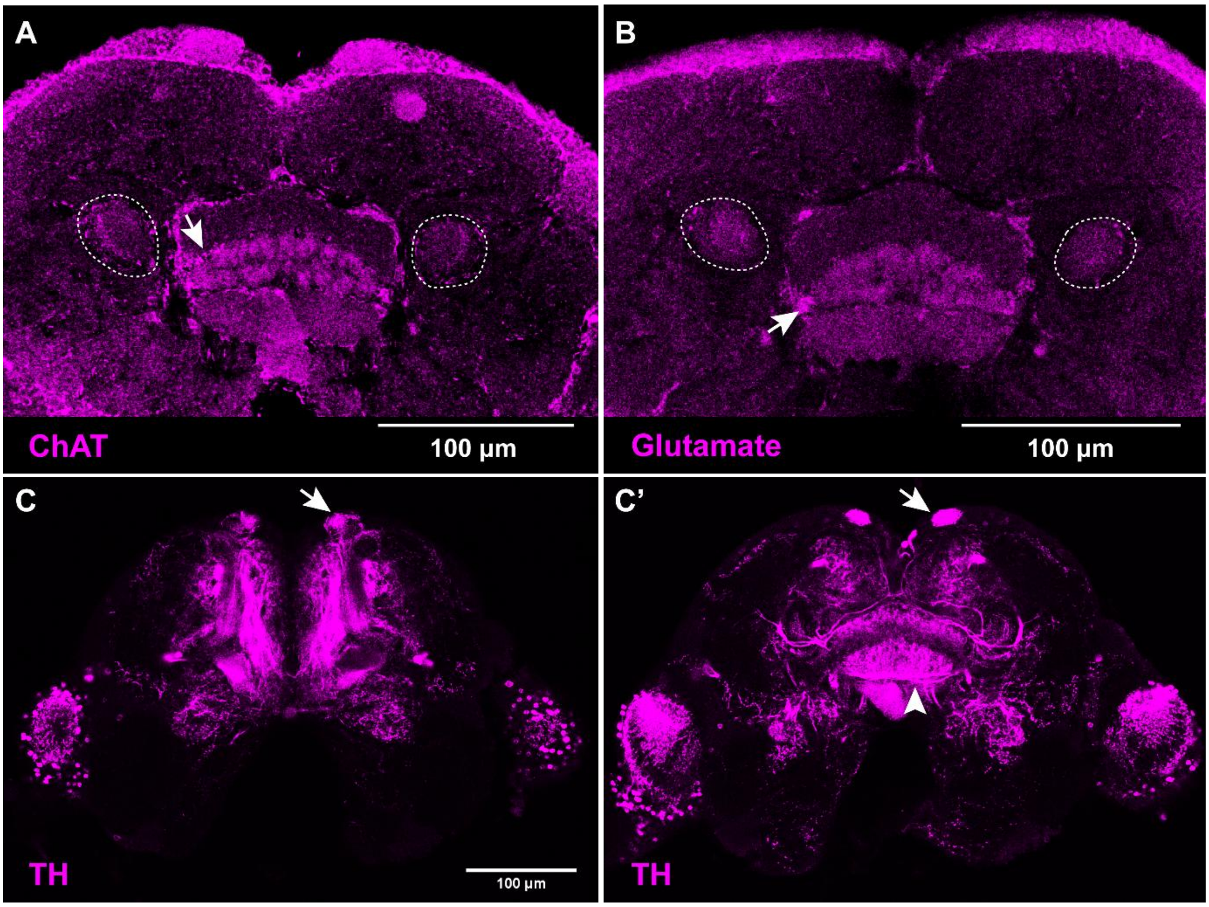


**Supplementary Figure 15.** Neurotransmitters expressed in the mushroom body and central complex. (A) Choline Acetyltransferase (ChAT). White dashed circles indicate the peduncle (PED) of the mushroom body. The white arrow indicates the signals in the central body. (B) Glutamate (GLUT). White dashed circles represent the peduncle (PED) of mushroom body. White arrow shows the signals in the central body. (C-C') Dopamine (TH). White arrows indicate the signals in the α-lobe of mushroom body. White arrowhead shows signals in the central body. Scale bar: 100 µm.

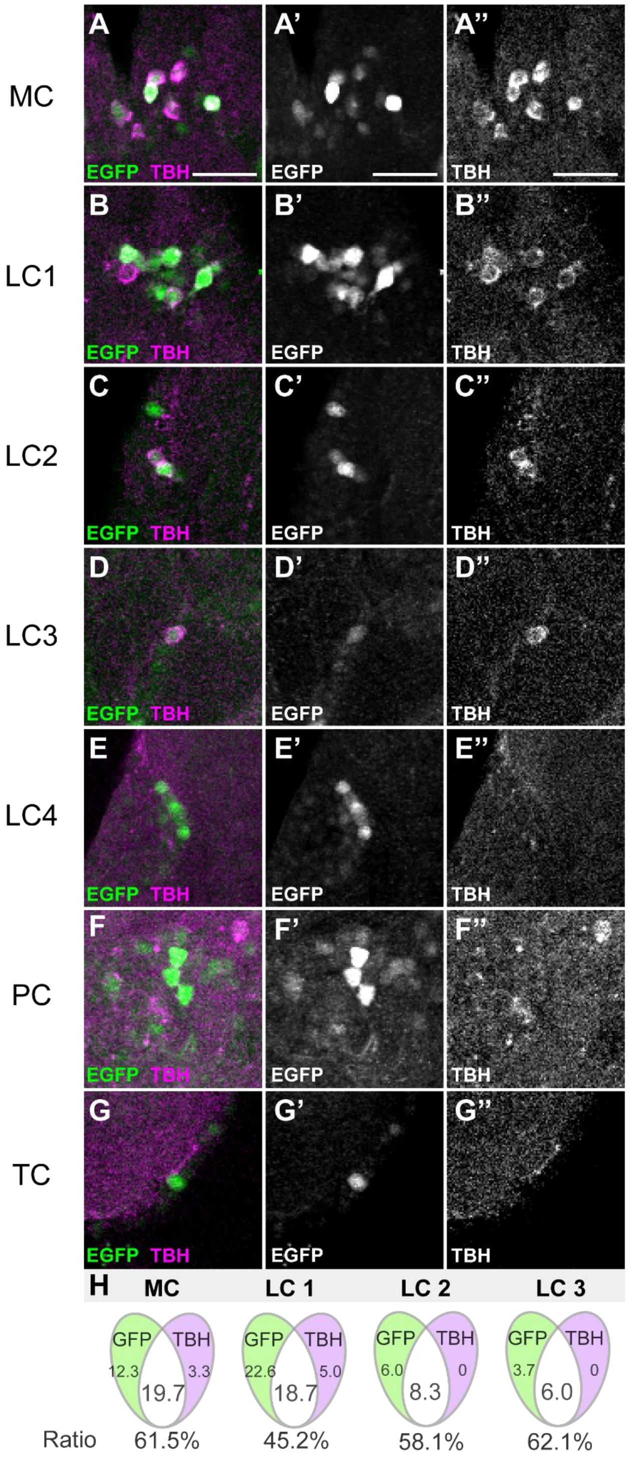


**Supplementary Figure 16.** Octopamine (TBH) is expressed in five FoxQ2-positive subclusters. (A-A'') The median cluster (MC), including MC1 and MC2, expresses both TBH and EGFP. (B-B'') In Lateral cluster 1 (LC1), TBH and EGFP largely overlap. (C-C'') In Lateral cluster 2 (LC2), TBH and EGFP largely overlap. (D-D'') Lateral cluster 3 (LC3) expresses both TBH and EGFP. (E-E'') Lateral cluster 4 (LC4) expresses only EGFP. (F-F'') In posterior cluster (PC), no overlap was detected between TBH and EGFP. (G-G'') The tritocerebrum cluster (TC) expresses only EGFP. (H) Venn diagrams illustrating the overlap between octopamine-positive and FoxQ2-positive neurons across the four subtypes (n = 3). Scale bar: 20 µm.

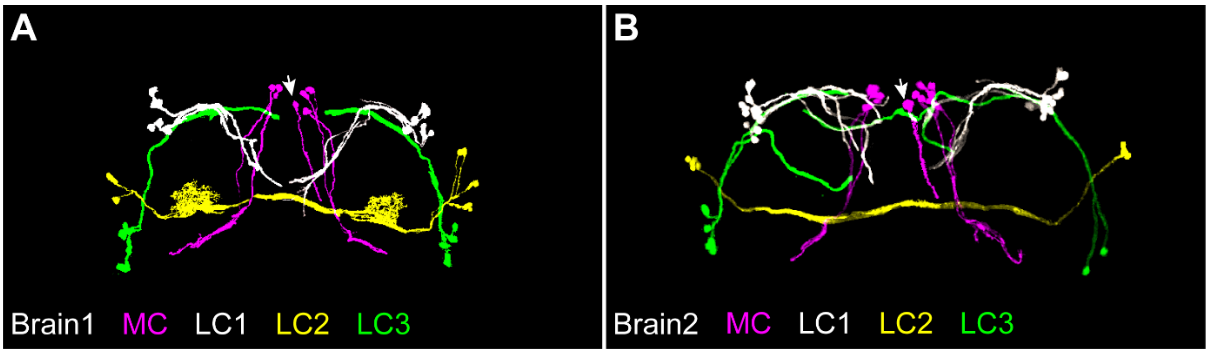


**Supplementary Figure 17. Reconstruction of dopamine/Tc-foxQ2 double positive neurons helps to confirm subclusters**. Dopaminergic neurons among EGFP-positive cells were reconstructed using the VVD viewer. (A-B) Overview images of two brains different from the one shown in Fig. 11. The median cluster (MC) is shown in magenta; lateral cluster 1 (LC1) in white; lateral cluster 2 (LC2) in yellow; and lateral cluster 3 (LC3) in green. In the MC, the white arrow indicates MC2, while the remaining region corresponds to MC1. (n=2). Only in the sample shown in A, the arborizations were clear enough to reconstruct them.

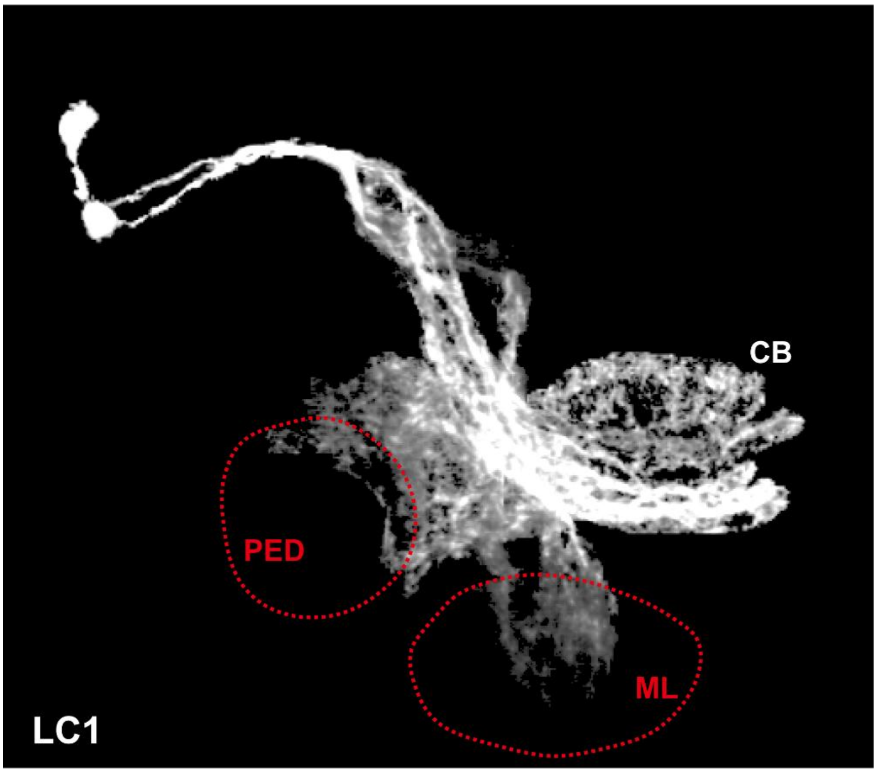


**Supplementary Figure 18.** *Tc-foxQ2*-Dopaminergic neurons of the lateral cluster 1 (LC1) have arborizations int mushroom body (MB) and the central body (CB). The arborizations could stem from different neurons but we cannot rule out that one neuron projects into both neuropils. Red dashed circles indicate components of the MB. PED: peduncle, ML: medial lobe.

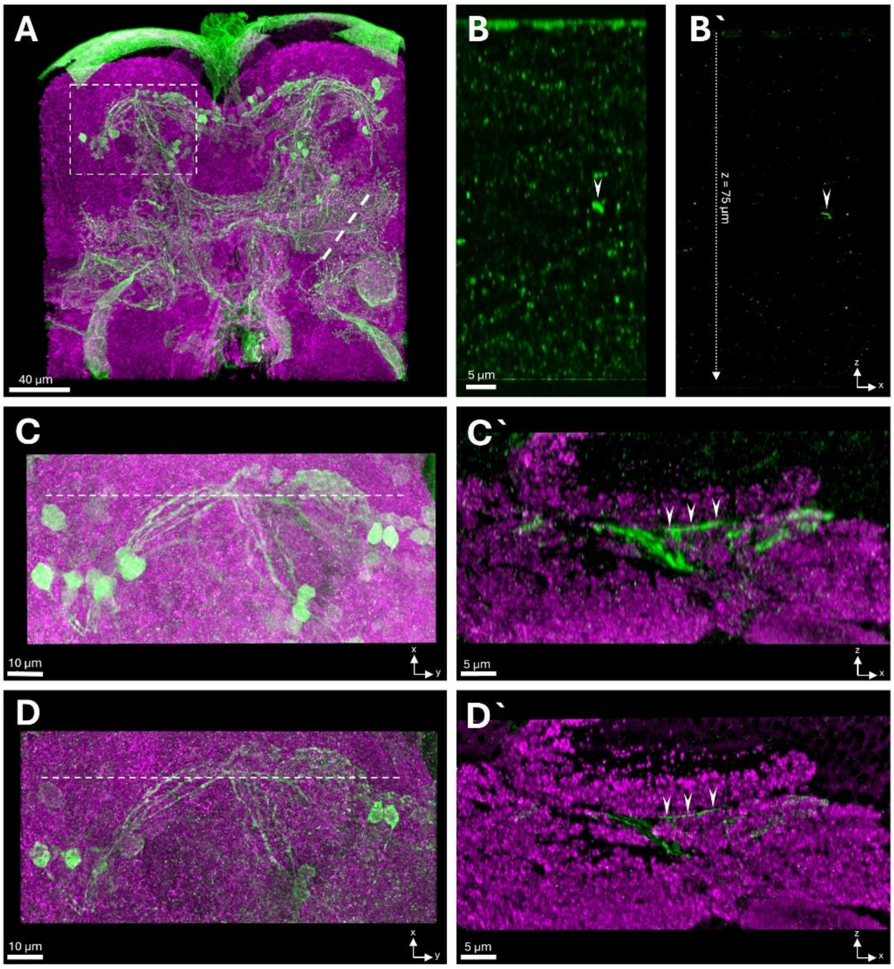


**Supplemenatary Figure 19. STED super-resolution microscopy in the *Tribolium* brain.** (A) Confocal overview of the central brain stained with Synapsin (magenta) and *Tc-foxQ2* positive neurons (green). Dashed line indicates imaging position in B, dashed rectangle shows imaging position in C and D. (B-B`) xz-scan (~75µm in z, FoxQ2 channel only) through the the brain in confocal (B) and 3D-STED (B`) mode, showing the resolution increase in 3D-STED. The white arrowhead shows a homogeneous structure in confocal mode, while in 3D-STED clearly 4 individual neurites can be distinguished. (C) Projection view of a confocal volume. Dashed white line indicates the position of the section in C`. (C`) Confocal xz-section. White arrowheads indicate an individual FoxQ2-positive neurite (green) running through the neuropil tissue (magenta, Synapsin staining). (D) Projection view of a STED (green, FoxQ2 positive neurons) and confocal volume (magenta, Synapsin, deconvolved). Dashed white line lindicates the position of the section in D`. (D`) STED xz-Section. White arrowheads indicate the same individual FoxQ2-positive neurite shown in C` running through the neuropil tissue (magenta, Synapsin). A comparison of the neurite in C` and D` clearly shows the resolution increase in STED compared to the confocal image.
